# The zinc-finger transcription factor LSL-1 is a major regulator of the germline transcriptional program in *C. elegans*

**DOI:** 10.1101/2021.12.16.472952

**Authors:** David Rodriguez-Crespo, Magali Nanchen, Shweta Rajopadhye, Chantal Wicky

**Affiliations:** Department of Biology, University of Fribourg, 1700 Fribourg, Switzerland

**Keywords:** *Caenorhabditis elegans*, chromatin, transcription regulation, RNA-seq, ChlP-seq, germline, meiosis

## Abstract

Specific gene transcriptional programs are required to ensure proper proliferation and differentiation processes underlying the production of specialized cells during development. Gene activity is mainly regulated by the concerted action of transcription factors and chromatin proteins. In the nematode *C. elegans*, mechanisms that silence improper transcriptional programs in germline and somatic cells have been well studied, however, how are tissue specific sets of genes turned on is less known. LSL-1 is herein defined as a novel crucial transcriptional regulator of germline genes in *C. elegans*. LSL-1 is first detected in the P4 blastomere and remains present at all stages of germline development, from primordial germ cell proliferation to the end of meiotic prophase. *lsl-1* loss-of-function mutants exhibit many defects including meiotic prophase progression delay, a high level of germline apoptosis, and production of almost no functional gametes. Transcriptomic analysis and ChIP-seq data show that LSL-1 binds to promoters and acts as a transcriptional activator of germline genes involved in various processes, including homologous chromosome pairing, recombination, and genome stability. Furthermore, we show that LSL-1 functions by antagonizing the action of the heterochromatin proteins HPL-2/HP1 and LET-418/Mi2 known to be involved in the repression of germline genes in somatic cells. Based on our results, we propose LSL-1 to be a major regulator of the germline transcriptional program during development.

## INTRODUCTION

Sexual reproduction relies on the generation of functional gametes, which depends on the proliferation and differentiation of primordial germ cells (PGCs) into oocytes and sperms. To ensure proper gametogenesis gene activity must be tightly regulated from the birth of PGCs to the production of mature sperm and oocytes including proper progression through meiosis. In every organism studied to date, gene regulation mechanisms represent an intrinsic part of the germ cell specification process (Seydoux and Braun 2006; Strome and Updike 2015). Studies across species have been mainly focused on how transcription of the somatic program is silenced and propose two modes of transcription repression in PGCs. Initially, transcription is blocked by the inhibition of transcription elongation at the level of RNA polymerase II, and eventually, a chromatin-based transcription repression takes over later in development (Nakamura and Seydoux 2008; Updike *et al*. 2014; Strome and Updike 2015; Seydoux 2018). In the *C. elegans* germline blastomeres, the PIE-1 protein sequesters the elongation factor P-TEFb. P-TEFb is a cyclin-dependent kinase that phosphorylates the CTD domain of polymerase II to allow transcription elongation (Ghosh and Seydoux 2008). Later, inhibition of the somatic transcriptional program switches to a chromatin-based repression. In *C. elegans* embryos, at about 100-cell stage, the P4 blastomere gives birth to PGCs Z2/Z3, PIE-1 disappears, and PGCs chromatin becomes depleted of di-methylated lysine 4 of histone H3 (H3K4me2)—a mark of active chromatin—and enriched in H3K9me—a mark of repressed chromatin (Seydoux and Dunn 1997; Schaner and Kelly 2006; Strome and Updike 2015). Loss of H3K4 methylation depends on the RNA binding proteins NOS-1 and NOS-2 (Schaner *et al*. 2003). However, PGCs Z2/Z3 are not completely transcriptionally silent. Zygotic expression of a few germline genes is detected: the P granules components (e.g., PGL-1), the germ cell fate maintenance RNA-binding protein NOS-1, or the chromatin-associated proteins XND-1 and OEF-1 (Kawasaki *et al*. 2004; Wang and Seydoux 2013; Mainpal *et al*. 2015; McManus and Reinke 2018). Although, the H3K36 methyltransferase MES-4 is known to confer transcriptional competence to germline genes, the mechanism by which transcription is initiated in PGCs is not yet well understood (Rechtsteiner *et al*. 2010). Following the onset of transcription, chromatin continues to assume a protective role which is mediated by MES-2/3/6 proteins—the worm PRC2 complex—repressing the somatic transcriptional program (Tursun *et al*. 2011; Patel *et al*. 2012). Finally, robust transcription is initiated when larvae start to feed after hatching of the embryo. At this stage, PGCs start to proliferate and later, at the larval stage L3, these enter into meiosis and differentiate into sperm (at larval stage L4) and oocytes (at adult stage). For most germline specific genes studied in adult worms—with the exception of those active during spermatogenesis—promoters are permissive for transcription in all germ cells; proper patterning of gene expression requires the 3’ untranslated region (3’UTR). Specialized proteins FBF-1/2, GLD-1, and MEX-3 were identified as crucial for the posttranscriptional regulation at the level of the 3’UTR of mRNAs (Merritt *et al*. 2008).

Here, we report the functional characterization of LSL-1, a novel key transcription regulator of germline genes. LSL-1 protein is first detected in the P4 blastomere and maintained in PGCs and developing germ cells in the gonad. Absence of LSL-1 activity leads to chromosome pairing defects, high levels of apoptosis, and a very low production of functional gametes. Based on our transcriptome profiling experiments and ChIP-seq (chromatin immunoprecipitation followed by sequencing) analysis of data available from modERN (branched from modENCODE project), we propose that LSL-1 acts as a direct transcriptional activator of germline genes involved in different aspects of germline development, including meiotic prophase progression and genome stability. Furthermore, we found that the sterility of *lsl-1* mutants is depending on the heterochromatin factors HPL-2/HP1 and LET-418/Mi2, involved in the silencing of germline gene transcription in somatic cells. Altogether, this lead us to propose that LSL-1 is an important player in the activation of the germline transcriptional program.

## MATERIALS AND METHODS

### Genetics

Worms were grown and maintained at 15 °C and 20 °C under standard conditions (Brenner 1974). Experiments were performed at 20 °C unless otherwise stated. *C. elegans* var. Bristol (N2) was used as wild type. Standard genetic crosses were made to generate double mutants using strains previously backcrossed to N2 at least four times. A list of all strains used in this study is provided in the Supplemental Methods section.

### Brood size, embryonic viability, and incidence of male analysis

Synchronized L4 hermphrodite worms were individually placed on NGM plates seeded with *E. coli* OP50 and then transferred to new plates every 24 h until laying stopped. Total number of laid eggs, hatched larvae, progeny which reached adulthood, and males were scored. Each scoring experiment was performed at the indicated temperature with mutant strains and wild-type strain N2 running in parallel. Data were pooled from multiple rounds of analyses, and average brood size, embryonic viability, and incidence of males were determined. Statistical significance was assessed using two-tailed Student’s *t*-test with Welch’s correction, *p*-value ≤ 0.05.

### Immunofluorescence

One-day adult hermaphrodite gonads were processed and immunostained as described by Phillips *et al*. 2009, with various modifications. Detailed protocol is included in the Supplemental Methods section.

Early-staged embryos were obtained from gravid hermaphrodite dissection and then processed as described in the Supplemental Methods immunofluorescence section. Embryos were also obtained by hypochlorite treatment of gravid hermaphrodites (50mM NaOH + 1.25% NaOCl) (Lewis and Fleming 1995), thereby allowing the acquisition of late embryonic stages. Synchronized populations of worms for each larval stage were also collected in distilled water and fixed in the same manner as the bleached embryos, using a modified protocol from Finney and Ruvkun 1990 (Miller and Shakes 1995; Bettinger *et al*. 1996). Samples were fixed with 2% formaldehyde in 1x modified Ruvkun fixation buffer (MRFB) and immediately frozen in liquid nitrogen. These were then thawed and incubated on ice for 30 min with occasional inversion and washed 3 × 10 min in PBST with DAPI added between the second and third washes. Finally, slides were mounted with Vectashield H-1000 antifade mounting medium (Vector Laboratories; Burlingame CA, USA), stored at 4 °C, and imaged. A list of antibodies used is available in the Supplemental Methods section.

### DAPI-staining cytological analysis

At least seven gonads of each genotype were stained with DAPI to determine the length extension of the mitotic to meiotic transition zone. Transition zone-like nuclei were identified based on their chromatin morphology and characteristic crescent shape. Length extension was measured in nuclei rows along the distal–proximal axis of the gonad according to Crittenden *et al*. 2006, defining its limits as the most distal and proximal rows where at least two nuclei exhibited the typical crescent shape.

At least 20 gonads of each genotype were stained with DAPI to quantify the number of DAPI-staining bodies in the diakinetic oocytes. The most proximal oocyte to the spermatheca (−1 oocyte) in each gonad was considered for the scoring. Slides were examined using a Nomarski and fluorescent Zeiss Axioplan 2 microscope to visualize the DAPI-stained bodies.

Data obtained from the different quantifications were pooled from multiple rounds of experiments in each cytological analysis category; statistical comparation between genotypes was assessed using two-tailed Student’s *t*-test with Welch’s correction, *p*-value ≤ 0.05.

### Fluorescence *in situ* hybridization (FISH)

A probe was generated from the 5S rDNA locus (located close to the pairing center region of chromosome V) by PCR (primer sequences) incorporating allyl-dUTP and labeled with the ATTO-488 NHS-ester fluorescent dye, as described in Sharma and Meister 2020. FISH probe hybridization was adapted from Phillips *et al*. 2009 and is described in detail in the Supplemental Methods section.

### Meiotic homologous chromosomes pairing dynamics analysis

To evaluate the progression of the chromosome pairing process, we monitored the localization of SUN-1::mRuby, HIM-8 (X-chromosome), and 5S rDNA probe (chromosome V) signals. At least four gonads of 24 h post-L4 hermaphrodite worms of each genotype were analyzed. Germlines were divided in seven equally long regions, from the distal tip to the proximal end of the pachytene stage, for a more precise comparison between wild-type and *lsl-1-mutant* strains.

Length extension of the germline region containing nuclei with the presence of SUN-1::mRuby signal patches was measured in nuclei rows along the distal–proximal axis. Quantification of HIM-8 and 5S rDNA foci involved scoring of the foci number observed per nucleus (n = 1: paired chromosomes; n > 1: unpaired chromosomes) in each germline region. Statistical comparisons were performed using two-tailed Student’s *t*-test with Welch’s correction, *p*-value ≤ 0.05.

### Microscopy and image processing

Imaging for the *lsl-1* expression pattern determination and germline cytological analysis was performed with a confocal microscope Leica TCS SPE-II DM5500Q. Images were collected using a 40x or 63x 1.3 NA objective (with 1.5x auxiliary magnification in embryos), and Z-stacks were set at 0.2 μm thickness intervals (0.5 μm for the cytological analysis of 1-day adult gonads). Embryos were staged by either morphology or number of blastomeres in early embryonic stages; larvae were staged by size or germline developmental phase from synchronized populations.

Images for meiotic chromosome pairing dynamics analysis were obtained using a pco.edge sCMOS camera attached to a Visitron Visiscope CSU-W1 spinning disk confocal microscope (Nikon Ti/E inverted microscope). Imaging was performed using a 100x 1.4 NA objective, and Z-stacks were set at 0.2 μm thickness intervals.

Total length of larval and 1-day adult gonads images were obtained as multiple Z-stacks due to their length and later merged to generate the complete final image using the ImageJ Stitching plugin (Preibisch et al. 2009) or Adobe Photoshop (2020). Images were processed using Fiji ImageJ, background was subtracted, and contrast/brightness adjusted. Orientation of the images and final figure appearance were performed using Adobe Photoshop (2020) and Adobe Illustrator (2020).

### Germline apoptosis

Apoptosis was determined using acridine orange (AO) staining in the germlines of 1-day-old hermaphrodite worms. Number of apoptotic corpses per gonad arm for wild-type and different single- and double-mutant strains was scored as in Shaham 2006. A detailed protocol is available in the Supplemental Methods.

### RNA extraction, cDNA library preparation, and sequencing

Wild type and both *lsl-1(tm4769)* and *lsl-1(ljm1)* mutant strains were synchronized and collected as young adult hermaphrodites 50-h postlarval hatching after hypochlorite treatment (at 20 °C). Total RNA was extracted with TRIzol Reagent (Invitrogen; Carlsbad CA, USA), and RNA was purified using the PureLink RNA Mini Kit (Invitrogen; Carlsbad IA, USA) according to manufacturer’s instructions. cDNA library preparation and RNA sequencing were performed at the Next Generation Sequencing (NGS) Platform in Bern (https://www.ngs.unibe.ch/). Quality and concentration of each RNA sample and following cDNA libraries were determined with Qubit 2.0 fluorometer and the Fragment Analyzer CE12 AATI. cDNA libraries were built using the TruSeq stranded mRNA library preparation kit (Illumina Inc.; San Diego CA, USA). RNA sequencing (50 bp paired-end reads) was performed on three biological replicates per sample with the Illumina NovaSeq 6000 Sequencing System, and cDNA libraries were multiplexed in a sequencing lane.

### RNA-seq data analysis

The sequencing data were obtained from Bern NGS platform. Raw reads in *.fasta* format were then uploaded to the Galaxy web platform using the public server at https://usegalaxy.org (Afgan *et al*. 2016). Sequencing data analysis is described in more detail in the Supplemental Methods.

### Chromatin immunoprecipitation (ChIP) sequencing data

We obtained the ChIP sequencing data analyzed in this study using the interface http://epic.gs.washington.edu/modERN/ that compiles the data from the model organism Encyclopedia of Regulatory Networks (modERN) consortium (Kudron *et al*. 2018)—branched from the model organism encyclopedia of DNA Elements (modENCODE) project (Gerstein *et al*. 2010)—available at: https://www.encodeproject.org/experiments/ENCSR969MNX/. ChIP-seq data processing and analysis are described in detail in the Supplemental Methods section.

## RESULTS

### *lsl-1* encodes a germ cell specific zinc-finger transcription factor

*lsl-1* (for *lsy-2*-like) was identified in a genome-wide RNAi screen as a suppressor of ectopic germline gene expression associated with mutations in *let-418*, which encodes an ATP-dependent chromatin remodeler (Erdelyi *et al*. 2017). *lsl-1* is predicted to encode a 318 aa protein with at least three zinc-finger domains (Figure 1A). This three-zinc-finger C2H2-type cluster is homolog to the zinc-finger domain which characterizes the SP/KLF family of transcription factors, a protein family with diverse functions in growth and development (Kaczynski *et al*. 2003; Pearson *et al*. 2008). In addition, two less conserved zinc fingers are located at the C terminal end of the protein (Figure S1). Along the entire length of the protein, LSL-1 shows 65% similarity to the LSL-1 paralog LSY-2—involved in ASE neuron specification and in the maintenance of germ–soma distinction—and 41% similarity to the zinc-finger domain of the human protein ZPF57, which plays a role in the allelic expression of imprinted genes (Figure S1 and Figure S2) (Alonso *et al*. 2004; Johnston 2005; Lin *et al*. 2015; Liu *et al*. 2017). The *Caenorhabditis elegans* Mutant Consortium 2012, provided a 675 bp deletion allele, *tm4769*, that removes the first three exons and part of the promoter region (Figure 1A, Table 1 and Table S1), and we generated an additional allele, *ljm1*, by inserting two consecutive stop codons 27 bp downstream the *lsl-1* translational initiation site (Figure 1A). *lsl-1(tm4769)* and *lsl-1(ljm1)* homozygous animals exhibit decreased brood size and embryo viability as well as a high incidence of males with respect to wild-type animals, indicating defects in meiotic prophase progression. Although both represent strong loss-of-function alleles, these exhibit a slightly different penetrance of the phenotype (Table 1). The present study was performed with the *tm4769* allele and is supplemented with data on the *ljm1* allele (Supplemental Material).

**Figure 1.**
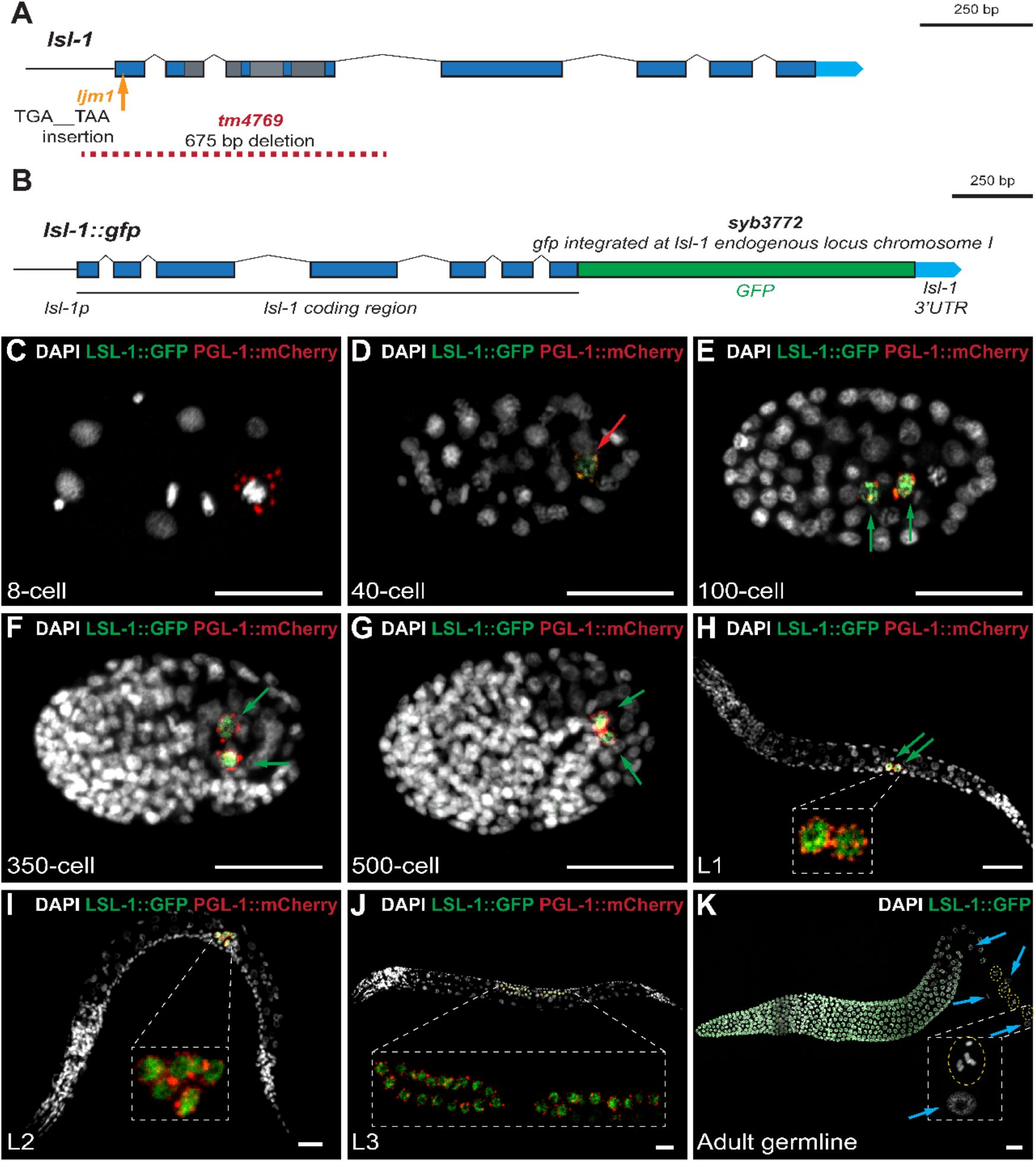
*lsl-1* is specifically expressed in the germline throughout development. (A) *lsl-1* gene structure and alleles used in this study. Blue boxes indicate exons. Light-blue arrows represent the 3’ UTR. Straight lines indicate promoters, and peaked lines, introns. Exon regions in grey encode the three Zinc-finger domains. (B) CRISPR/Cas9 mediated knock-in of the GFP coding sequence (green box) at the endogenous *lsl-1* gene locus (*syb3772[lsl-l::GFP]*). Representative confocal projection images of (C–J) whole worms at different developmental stages and (K) a dissected adult hermaphrodite gonad. Chromatin is stained with DAPI (grey), PGL-1::mCherry is in red, and LSL-1::GFP, in green. Red arrow denotes P4 blastomere; green arrows, primordial germ cells Z2/Z3; and light-blue arrows, somatic cell nuclei of the adult gonad (sheath cells). Dashed yellow ovals mark the diakinetic oocyte nuclei. Scalebars, 20 μm.

**Table 1.** Brood size, survival rate, and incidence of male (20 °C)

| Genotype | Mean brood size<sup>a</sup> | Viability (%) | Incidence of males (%) | *n*<sup>b</sup> |
| --- | --- | --- | --- | --- |
| wild type | 301.79 ± 31.25 | 98.44 | 0.06 | 37 |
| *lsl-1(tm4769)* | 28.10 ± 24.14*** | 0.12 | n/a<sup>c</sup> | 58 |
| *lsl-1(ljm1)* | 57.89 ± 27.18*** | 5.44 | 19.31 | 47 |
| *lsl-1(syb3772[lsl-1::GFP])* | 298.17 ± 78.93<sup>n.s</sup> | 98.83 | 0.08 | 12 |

<sup>a</sup> Data correspond to the mean ± SD of the total number of eggs laid per hermaphrodite parent. Statistical comparison between wild type and each genotype performed by two-tailed Student's *t*-test with Welch's correction. \*\*\* *p*-value ≤ 0.001, <sup>n.s</sup> *p*-value > 0.05.
<sup>b</sup> Total number of parental hermaphrodites per genotype.
<sup>c</sup> Not applicable (n/a)

The mutant phenotype suggests a function of LSL-1 in the germline. Using a LSL-1::GFP endogenous reporter, we examined the *lsl-1* expression pattern throughout development (Figure 1B). CRISPR/Cas9 mediated knock-in of the GFP coding sequence in the endogenous *lsl-1* gene upstream of the stop codon did not interfere with the protein function (Table 1). LSL-1 is detected in cells marked by the presence of P-granules, namely in the P4 blastomere and later on in primordial germ cells Z2 and Z3 throughout embryogenesis (Figure 1, C–G). During larval development and adult stage, LSL-1 is observed in proliferative germline nuclei and in pachytene and diplotene stage nuclei (Figure 1, J and K). LSL-1::GFP signal disappears at the late diplotene stage and is barely detectable in oocytes and sperm (data not shown) (Figure 1K). LSL-1 was not detected in somatic cells, including the gonadal sheath cells and the distal tip cell (Figure 1K). These results indicate that *lsl-1* is specifically expressed in germ cells throughout development and is essential for the production of functional gametes.

### LSL-1 is essential for normal progression of germ cells through meiotic prophase

To further investigate the function of LSL-1 in the germ cells, we performed a cytological analysis of *lsl-1* mutant gonads using DAPI staining. This analysis reveals the progression of nuclei through the different stages of the meiotic prophase based on chromatin organization (Figure 2A). By scoring the number of nuclei rows along the distal– proximal axis of the gonad, according to Crittenden *et al*. 2006, we observed an extended transition zone in *lsl-1(tm4769)* worms (32.9 ± 4.5 rows), with respect to the wild-type transition zone, comprised of 14.4 ± 20 rows in average (Figure 2, A and B). Similar results were observed in *lsl-1(ljm1)* mutant gonads (Figure S3). These observations suggest that chromosome pairing might be perturbed in *lsl-1* mutants.

**Figure 2.**
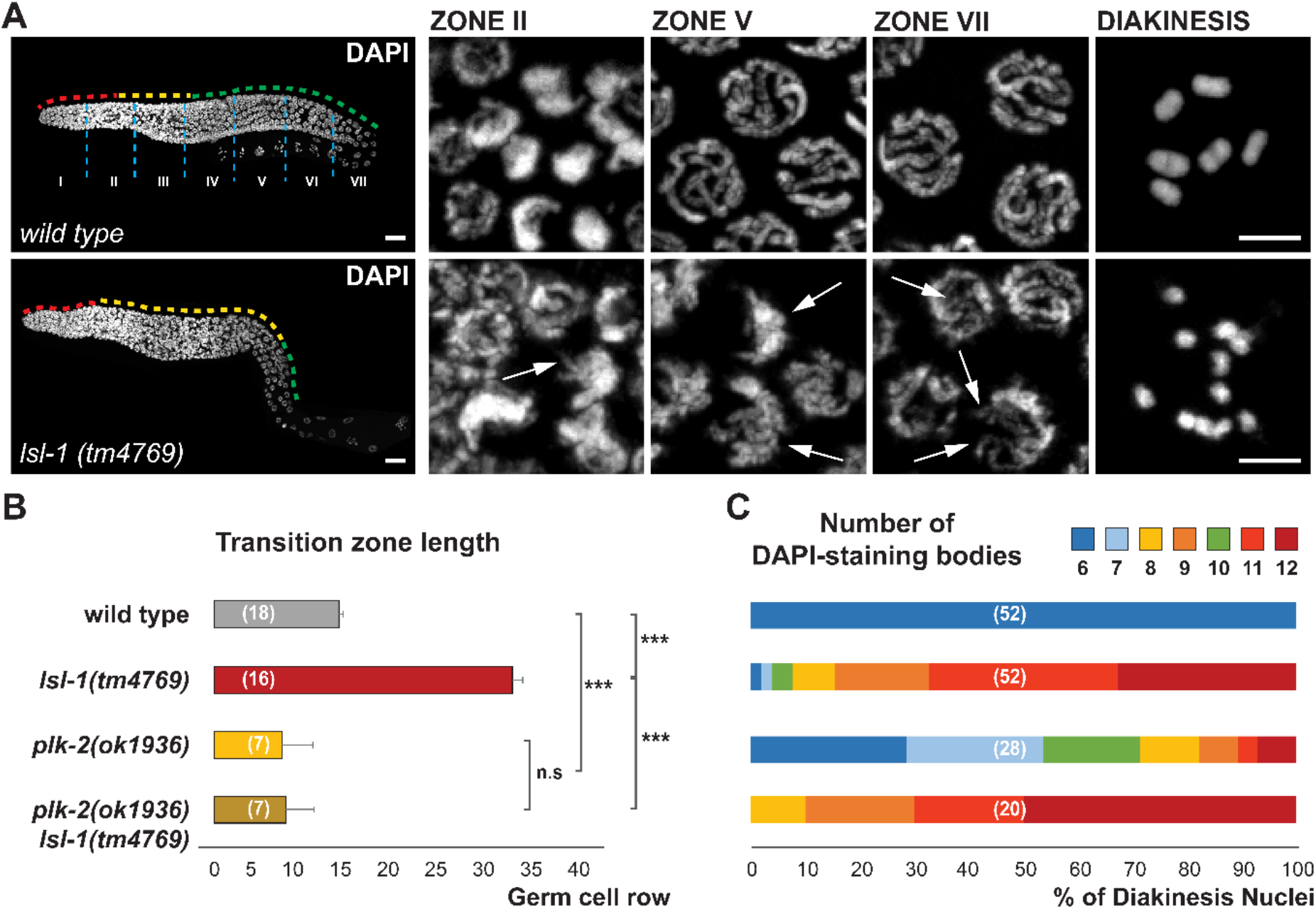
*lsl-1* mutants exhibit an extended transition zone that depends on PLK-2 and an altered chromatin organization in meiotic nuclei. (A) Representative confocal projection images of DAPI-stained gonads at 1-day-old adult stage of the indicated genotype. Each panel shows a magnification of the indicated zones. Dashed lines depict the mitotic region (red), the transition zone to meiosis (yellow), the pachytene stage (green), and the seven equally long zones in which gonads have been divided (light blue). Arrows point to altered chromatin structures (see text). At least 15 gonads were analyzed for the indicated genotypes. (B) Graphic representation of the transition zone length quantified in nuclei rows from the MR/TZ boundary to the TZ/PS limit. Data are plotted as horizontal bars that represent mean length. Error bars correspond to standard error (SEM). *p*-value ≤ 0.001 (***); *p*-value > 0.05 nonsignificant (n.s), by two-tailed Student’s *t*-test with Welch’s correction. Number of germlines scored for each genotype in brackets. (C) Percentages of diakinetic oocytes by number of DAPI-staining bodies content in 1-day-old adult hermaphrodite germlines for the indicated genotypes. Note, DAPI-staining bodies were scored from the oocytes immediately prior to spermatheca entry. Number of oocytes scored for each genotype in brackets. Scalebars, 20 μm and 5 μm in whole gonad images and magnification panels, respectively. MR, mitotic region; TZ, transition zone; PS, pachytene stage.

Closer examination of the DAPI-stained nuclei in defined zones of equal size along the gonad of *lsl-1* mutants revealed an altered chromatin organization in comparison with wild type. In the transition zone of *lsl-1* mutants, chromatin appears to loop out of the otherwise normally clustered chromosomes (Figure 2A, zone II and Figure S3 A, zone II). The few pachytene stage nuclei in *lsl-1* mutants exhibit disorganized chromosomes, with thinner stretches of chromatin which could represent unpaired regions of the chromosomes (Figure 2A, zones V–VII and Figure S3 A, zones V–VII).

In *lsl-1(tm4769)* mutant allele, we found less than 1.8% of *lsl-1* oocytes presenting the normal six DAPI-staining bodies, while the remaining 98.2% showed more than six DAPI-staining bodies (90% in *lsl-1(ljm1)* mutant allele) (Figure 2C and Figure S3 C). This indicates that a large portion of the chromosomes fail to undergo crossing over. These cytological defects are consistent with the high incidence of males and the decreased embryo viability observed in the progeny of *lsl-1* mutants, which likely result from chromosome missegregation at meiotic division I (Table 1 and Table S1).

### PLK-2 dependent cell cycle delay is activated in *lsl-1* mutants

The Polo-like kinase PLK-2 coordinates cell cycle delay and chromosome pairing (Fridkin *et al*. 2009; Harper *et al*. 2011). To determine whether the *lsl-1* extended transition zone depends on PLK-2 activity, we generated a *plk-2(ok1936) lsl-1(tm4769)* double mutant and scored the length of their transition zone. The double mutants *plk-2(ok1936) lsl-1(tm4769)* exhibit a transition zone length comparable to *plk-2(ok1936)* mutants, which is shorter than the wild-type transition zone (Figure 2B). *plk-2(ok1936) lsl-1(tm4769)* worms are sterile and present a slightly higher number of univalents compared to *lsl-1(tm4769)* mutants (Table 2, Figure 2C). These results suggest that PLK-2-dependent cell cycle delay is activated in the *lsl-1* mutant and might allow some level of pairing and recombination.

**Table 2.** Brood size, survival rate, and incidence of males (25 °C)

| Genotype | Mean brood size<sup>a</sup> | Viability (%) | Incidence of males (%) | *n*<sup>b</sup> |
| --- | --- | --- | --- | --- |
| wild type | 196.78 ± 42.22 | 94.46 | 0.09 | 46 |
| *lsl-1(tm4769)* | 0.05 ± 0.221 | 0.00 | n/a | 40 |
| *plk-2(ok1936)* | 83.82 ± 35.42 | 16.81 | 21.94 | 11 |
| *plk-2(ok1936) lsl-1(tm4769)* | 0.00 ± 0.00<sup>n.s</sup> | n/a | n/a | 12 |

<sup>a</sup> Data correspond to the mean ± SD of the total number of eggs laid per hermaphrodite parent. Statistical comparison between *plk-2(ok1936) lsl-1(tm4769)* double mutant and *lsl-1(tm4769)* genotype performed by two-tailed Student's *t*-test with Welch's correction. <sup>n.s</sup> *p*-value > 0.05.
<sup>b</sup> Total number of parental hermaphrodites per genotype.
n/a not applicable

### Chromosome pairing is disrupted in absence of LSL-1

To test whether homologous chromosome pairing is perturbed in *lsl-1* mutants, we monitored the localization of SUN-1, which forms aggregates upon phosphorylation by checkpoint kinase CHK-2 and polo-like kinase PLK-2, following pairing initiation (Fridkin *et al*. 2009; Woglar *et al*. 2013). Using SUN-1::mRuby transgenic worms, we observed SUN-1 aggregates at the beginning of the transition zone in *lsl-1* and wild-type gonads (Figure 3 and Figure S4). However, in *lsl-1* mutants, SUN-1 patches were still detectable at the most proximal part of the germline, as far as zone VI, where no SUN-1 patches iare detected in wild type (Figure 3 and Figure S4, zone VI), supporting the idea that LSL-1 is involved in the proper progression of the pairing process.

**Figure 3.**
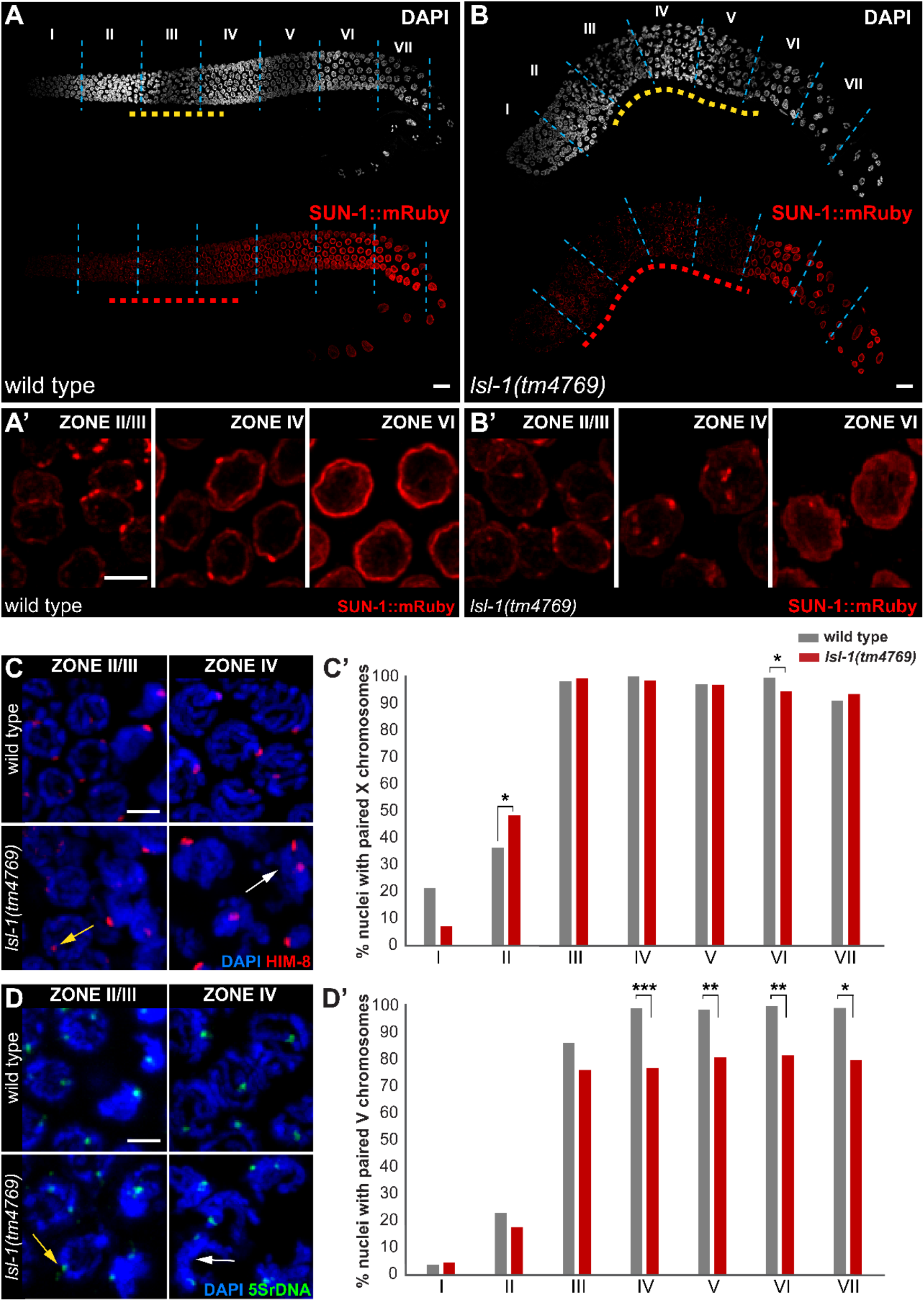
LSL-1 is required for the proper progression of homologous chromosome pairing. (A and B) Representative confocal projection images of 1-day-old adult stage gonads of (A) wild type and (B) *lsl-1(tm4769)* animals, expressing SUN-1::mRuby (red) and stained with DAPI (grey). Dashed lines delineate nuclei showing SUN-1::mRuby patches (red), extension of the transition zone (yellow), and the boundaries between the seven equally long zones (light blue). (A’ and B’) Each panel represents a magnification of the indicated zones. (C and D) Representative images of zone II/III and zone IV nuclei of the indicated genotypes: (C) immunostained with HIM-8 antibody (red); (D) hybridized with 5S rDNA FISH probe (green) to monitor chromosome pairing and costained with DAPI (blue). Arrows point to possible precocious paired chromosomes in late mitotic zone (yellow) or nuclei with unpaired signals (white). (C’ and D’) Histograms showing the percentage of nuclei with paired (C’) HIM-8 and (D’) 5S rDNA signals, scored per zones of the indicated genotypes. *p*-value ≤ 0.001 (***); *p*-value ≤ 0.01 (**); *p-* value ≤ 0.05 (*); *p*-value > 0.05 nonsignificant, by two-tailed Student’s *t*-test with Welch’s correction. At least three gonads from independent experiments were scored for each genotype. Scale bars, 20 μm and 5 μm in whole gonad images and magnification panels, respectively.

To better identify the pairing defects in *lsl-1* mutants, we monitored the localization of the X-chromosome Pairing Center (PC) protein HIM-8 using immunofluorescence (Figure 3C and Figure S4). Paired chromosomes present one HIM-8 focus per nucleus. In wild-type animals, more than 90% of X chromosomes are paired from zone III to the most proximal regions of the germline (Figure 3C’ and Figure S4 C’). In *lsl-1(tm4769)* mutants, an increased number of single HIM-8 per nucleus is observed, suggesting that precocious X-chromosome pairing could occur (Figure 3C’ and Figure S4 C’; zone II *p*-value ≤ 0.05). However, this observation could also be due to a slightly shorter mitotic zone in *lsl-1* mutants (Figure S5). Although the overall number of nuclei in the mitotic zone is not significantly different between wild type and *lsl-1* mutants, we observe a shorter mitotic region and a decreased mitotic index in *lsl-1* mutants compared with wild type (Figure S5). Zones III to VII exhibit a similar level of pairing in *lsl-1* mutants and wild-type worms; however, a significant decrease in the number of single HIM-8 foci is observed in *lsl-1(tm4769)* zone VI in comparison with wild-type gonads indicating minor perturbations in the pairing process (Figure 3C; *p*-value ≤ 0.05).

Pairing of chromosome V was investigated using a FISH probe made of 5S rDNA repeats. This approach showed that pairing of chromosome V never reaches wild-type level in *lsl-1* mutants (Figure 3, D and D’; Figure S4, D and D’). From zone IV up to zone VII, a significant decrease is observed in the level of pairing (Figure 3D’ and Figure S4 D’). These observations indicate that chromosome pairing is compromised in *lsl-1* mutants, however, not to an extent that could explain the high number of univalents observed at diakinesis.

### Absence of LSL-1 activity triggers elevated apoptosis levels

*lsl-1* mutants lay a very limited number of embryos, suggesting that a high number of germline nuclei might be eliminated by apoptosis. Using acridine orange staining, we observed a significant increase in the number of apoptotic germ cells in *lsl-1* mutants compared to wild-type worms (Figure 4 and Figure S6). This elevated number of apoptosis could be due to the activation of pairing and/or DNA damage checkpoints (Harper *et al*. 2011; Kim *et al*. 2015; Mateo *et al*. 2016). To test pairing checkpoint activation, we measured the level of apoptosis in *lsl-1* mutant germlines lacking *plk-2* activity. *plk-2(ok1936) lsl-1(tm4769)* double mutants show a decreased level of apoptosis compared to *lsl-1* mutants, more similar to wild-type apoptosis level. This result is consistent with pairing defects triggering PLK-2 dependent apoptosis.

**Figure 4.**
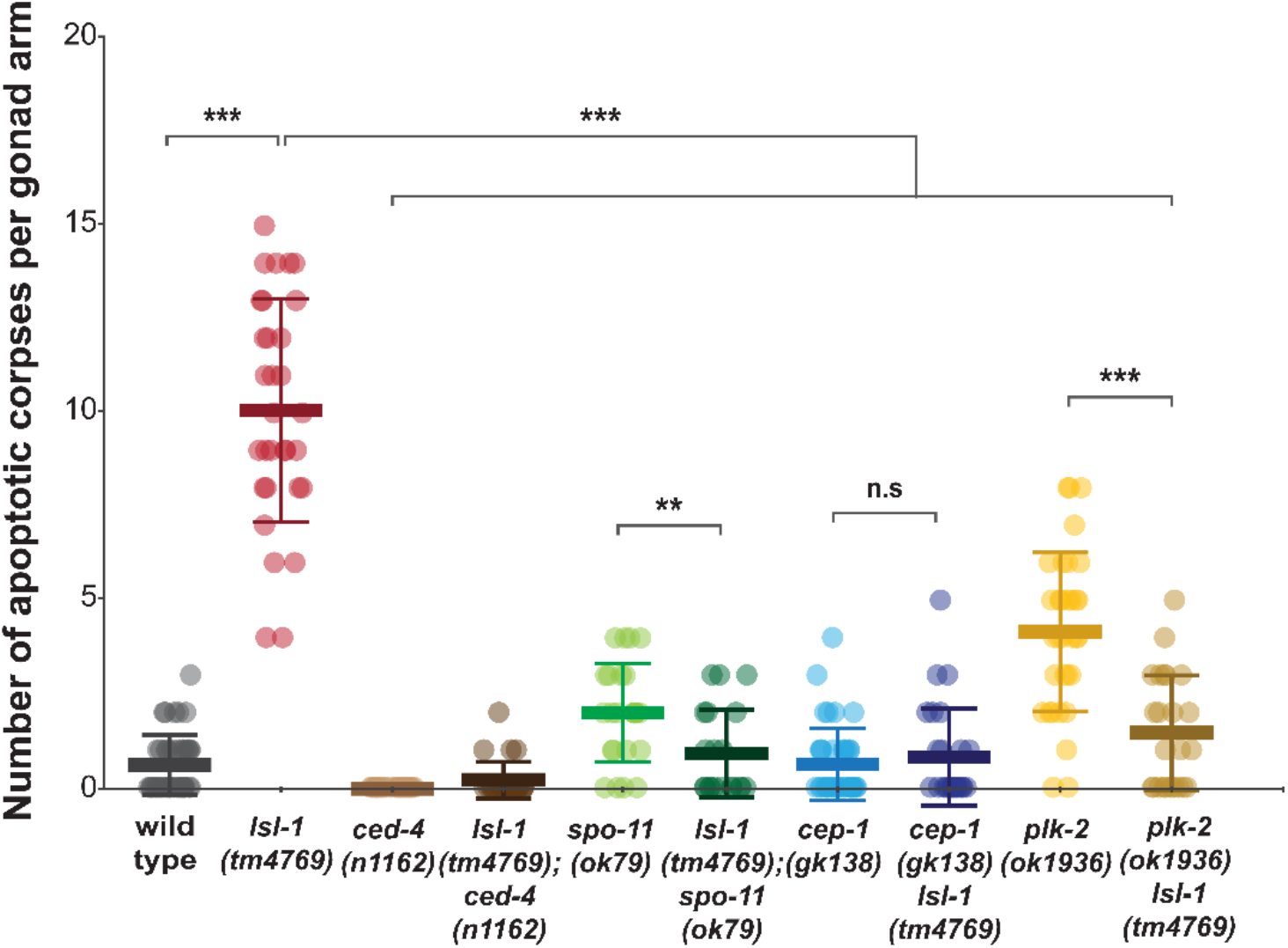
Increased level of apoptosis in *lsl-1* mutants is dependent on DNA damage and pairing checkpoints, CEP-1 and PLK-2, respectively. Scatter plot showing the number of apoptotic corpses per gonad arm from 1-day adult hermaphrodites for the indicated genotypes and quantified by acridine orange staining. Data are plotted as vertical dot plots, with each dot representing the number of apoptotic corpses in one gonad arm. Horizontal lines represent mean, with error bars corresponding to standard deviation (SD). *p*-value ≤ 0.001 (***); *p*-value ≤ 0.01 (**). *p*-value ≤ 0.05 (*); *p*-value > 0.05 nonsignificant (n.s), by two-tailed Student’s *t*-test with Welch’s correction. At least 24 gonads from different biological replicates were scored for each genotype.

In addition, *lsl-1* mutants could also exhibit an elevated apoptosis level due to persistent DNA damage and activation of the checkpoint protein CEP-1/p53 (Kim *et al*. 2015; Mateo *et al*. 2016). Absence of CEP-1 activity reduces the level of apoptosis in *cep-1(gk138) lsl-1(tm4769)* mutants to wild-type levels, indicating accumulation of DNA damage in the absence of LSL-1. Furthermore, in *lsl-1(tm4769); spo-11(ok79)* double mutant, which does not initiate recombination (Dernburg *et al*. 1998), the increased level of apoptosis is reduced to wild-type levels, indicating that persistent DNA damage results from unresolved recombination intermediates in *lsl-1(tm4769)* animals (Figure 4). Our overall results show that both synapsis defects and unresolved recombination events contribute to the high level of apoptosis observed in *lsl-1* mutants.

### LSL-1 regulates transcription of germline genes

Lsl-1 encodes a zinc-finger containing protein, which most functions as a transcriptional regulator. To test this, we performed a transcriptome RNA-seq analysis. A total of 978 upregulated and 1100 downregulated genes were identified in *lsl-1(tm4769)* mutants compared with wild-type animals (*q*-value ≤ 0.01; −2 ≥ fold change ≥ 2) (Figure 5A and File S1). Tissue enrichment analysis (Angeles-Albores *et al*. 2016) showed that the vast majority of downregulated genes were associated with germline functions and the reproductive system. In addition, genes specific to male functions, to neurons, and to the epithelial system were also found to be deregulated (Figure 5B and File S2). Among the genes involved in germline functions, we identified genes involved in germ cell fate (*nos-2, xnd-1*), in pairing and synapsis (*pch-2, sun-1, syp-2, zim-1, zim-3*), in genome stability (*chk-1, dsb-2, hsr-9*), in P granules composition (*glh-2, meg-4, mex-3, mex-6, oma-1, pgl-2, pie-1, pos-1*), or in the mitotic/meiotic transition (*fbf-1, glp-1*). Overall, these data indicate that LSL-1 regulates genes involved in several germline processes.

**Figure 5.**
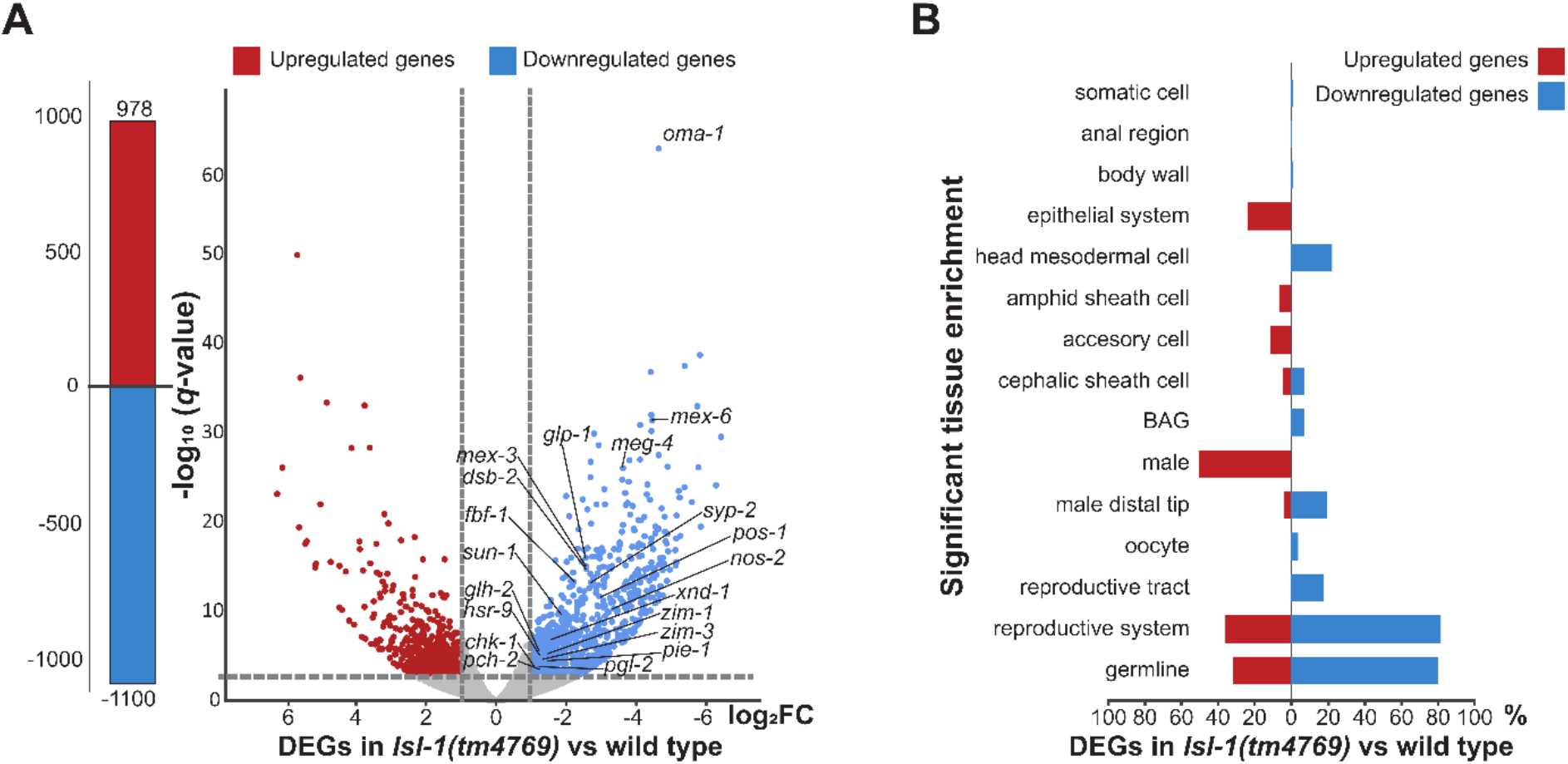
Absence of LSL-1 leads to germline gene expression changes. (A) Bar and volcano plots show the number of significant differentially expressed genes (DEGs) in *lsl-1(tm4769)* young adult animals compared with wild type determined by RNA-seq analysis. Each dot represents a gene, and red and blue colors correspond to significant up- and downregulated genes, respectively. Dash lines indicate the significance and fold change cutoffs (*q*-value ≤ 0.01 and −2 ≥ fold change ≥ 2). In italics, representative genes associated to different germline functions (see results section). Note, symbol (-) stands for downregulation, both in number of genes and fold change; *q*-value stands for adjusted *p*-values found using an optimized false discovery rate (FDR) approach (Storey and Tibshirani 2003). (B) Graph illustrating the tissue enrichment of significant up- (red) and downregulated (blue) genes in *Isl-1 (tm4769)* young adult animals compared with wild type, using the T.E.A-Wormbase tool (Angeles-Albores *et al*. 2016) and represented as a percentage of total significant up- or downregulated genes. Only enrichments with significant adj. *p*-value ≤ 0.05 were scored. For enrichment in blastomeres see File S2. FC, fold change; DEGs, differentially expressed genes; T.E.A, tissue enrichment analysis; BAG, neuron class of two neurons with ciliated endings, in the head, with elliptical, closed, sheet-like processes near the cilium, which envelop a piece of hypodermis (see Wormbase anatomy term).

The total number of DEGs in the *lsl-1(ljm1)* was lower (*n* = 496) than in the *lsl-1(tm4769)* allele (*n* = 2078) (Figure S7 A and File S1). However, 80% overlapped with the DEGs detected in *lsl-1(tm4769)* mutants (Figure S7 B). Common DEGs appeared deregulated in the same direction (File S3) and exhibited similar tissue enrichment patterns (Figure 5B, Figure S7 C and File S2). These results are consistent with the difference in penetrance observed in the phenotypes associated with the two alleles.

### LSL-1 binding sites are highly enriched on autosomes

To identify LSL-1 binding site to the genome, we analyzed ChIP-seq data available from the modERN consortium (Kudron *et al*. 2018). ChIP-seq was performed in worms carrying an LSL-1::TY1::EGFP::3xFLAG (*wgIs720*) transgene, whose expression matches our own observations (Figure S8).

A total of 3896 significant peaks were identified as enriched by the ChIP-seq processing pipeline (SPP) (Kharchenko *et al*. 2008), IDR < 0.1%, and mapped corresponding to 3078 genes in the *C. elegans* reference genome (version WS245) (File S4). Distribution of peaks did not reveal marked intrachromosomal bias (Figure 6A). However, LSL-1 is almost completely absent from the X-chromosome (*n* = 60) while highly enriched on chromosomes III and I (*n* = 901 and *n* = 869, respectively) (Figure 6B). This distribution pattern resembles the chromosomal distribution of germline-specific genes (Reinke and Cutter 2009; Rechtsteiner *et al*. 2010; Kelly *et al*. 2014). Peaks were narrow, with an average size of 400 bp (Figure 6C), and 74% of them were preferentially associated with promoters (defined as 2 kbp upstream of a gene) (Figure 6, D and E). Moreover, using MEME-ChIP tool (Machanick and Bailey 2011), we performed a motif enrichment analysis for the significant peaks, detecting a very significant enrichment for the motif TAC_GTA (Figure 6F and Figure S9). This corresponds to the motif described previously in protein binding microarray studies as a highly enriched motif in the upstream regions of the germline precursors associated genes (Narasimhan *et al*. 2015). Location of LSL-1 binding sites in gene promoters, chromosomal bias, and motif enrichment in promoters of precursor germ cells associated-genes together, reinforce the idea that LSL-1 could function as a transcriptional regulator of germline genes.

**Figure 6.**
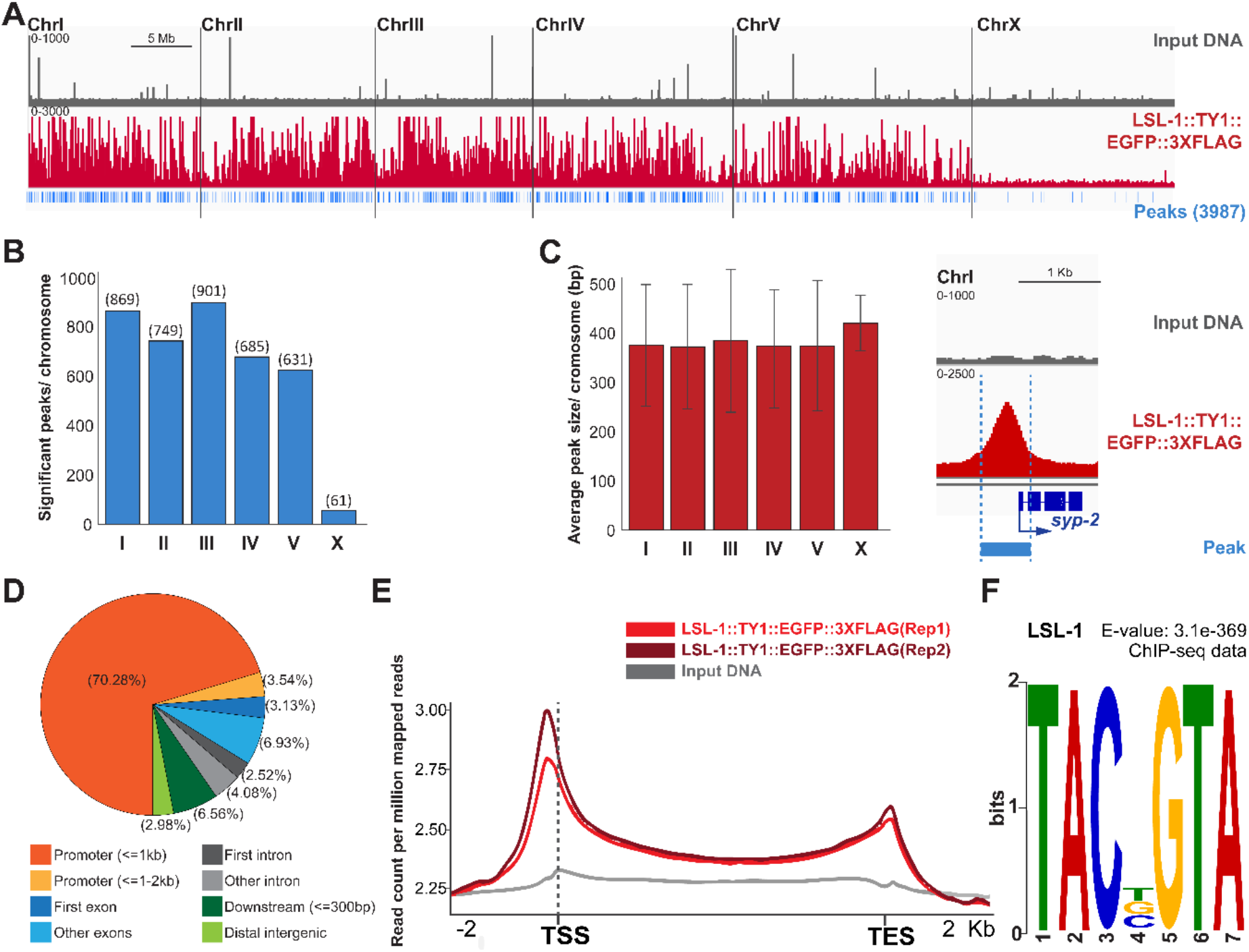
LSL-1 preferentially binds to promoters of genes localized on autosomes. (A) Genome-wide ChIP-seq binding profile of LSL-1::TY1::EGFP::3XFLAG (red line) compared with nonenriched input DNA (gray). Blue vertical bars (peaks) represent the most significant LSL-1 enriched regions (IDR ≤ 0.1%). (B) Graphic description of LSL-1::TY1::EGFP::3XFLAG significant ChIP-seq peaks distributed by chromosome (number of peaks in brackets). (C) Histogram illustrating the peak size per chromosome and a representative LSL-1::TY1::EGFP::3XFLAG ChIP-seq read pileup size for *syp-2* locus. Data are plotted as vertical bars that represent mean peak size. Error bars correspond to standard deviation (SD). (D) Graph shows the distribution of LSL-1 binding sites at the indicated genomic regions, in percentage. (E) Ngsplot of LSL-1 genome-wide enrichment centered on the gene body between TSS and TES. Input DNA is represented in gray for comparison, and results are represented as read counts per million reads. LSL-1 is mainly localized in gene promoter regions with a maximum peak upstream the TSS. (F) Illustration represents the most significant motif enriched in LSL-1::TY1::EGFP::3XFLAG peaks, identified using the MEME-ChIP platform (Machanick and Bailey 2011). The sequencing files of the LSL-1 ChIP-seq experiment analyzed here were performed in two different biological replicates and obtained from modERN consortium (Kudron *et al*. 2018), available at https://www.encodeproject.org/experiments/ENCSR969MNX/. TSS, transcription start site; TES, transcription end site; IDR, irreproducible discovery rate.

### LSL-1 is a transcriptional activator of germline genes

To identify LSL-1 direct target genes, we cross-compared our RNA-seq data and the ChIP-seq analysis. This overlap was significant, and the genes bound by LSL-1 represented 19% of the DEGs in *lsl-1(tm4769)* mutant (Figure 7A and Table S2). Remarkably, most of these genes were downregulated in *lsl-1* and mainly targeted at promoter regions, which suggests that LSL-1 acts as a transcriptional activator (Figure 7B and File S5). In addition, we performed a functional GO analysis using DAVID bioinformatics resources 6.8, NIAID/NIH tool with these potential LSL-1 direct targets (Huang *et al*. 2009). *lsl-1(tm4769)* gene set was significantly enriched in GO terms (*p*-value ≤ 0.05), such as embryo development, P granules, 3’ UTR-mediated mRNA destabilization, germ cell development, or meiotic division (Figure 7B’); similar results were obtained from the *lsl-1(ljm1)* dataset (Figure S10 C).

**Figure 7.**
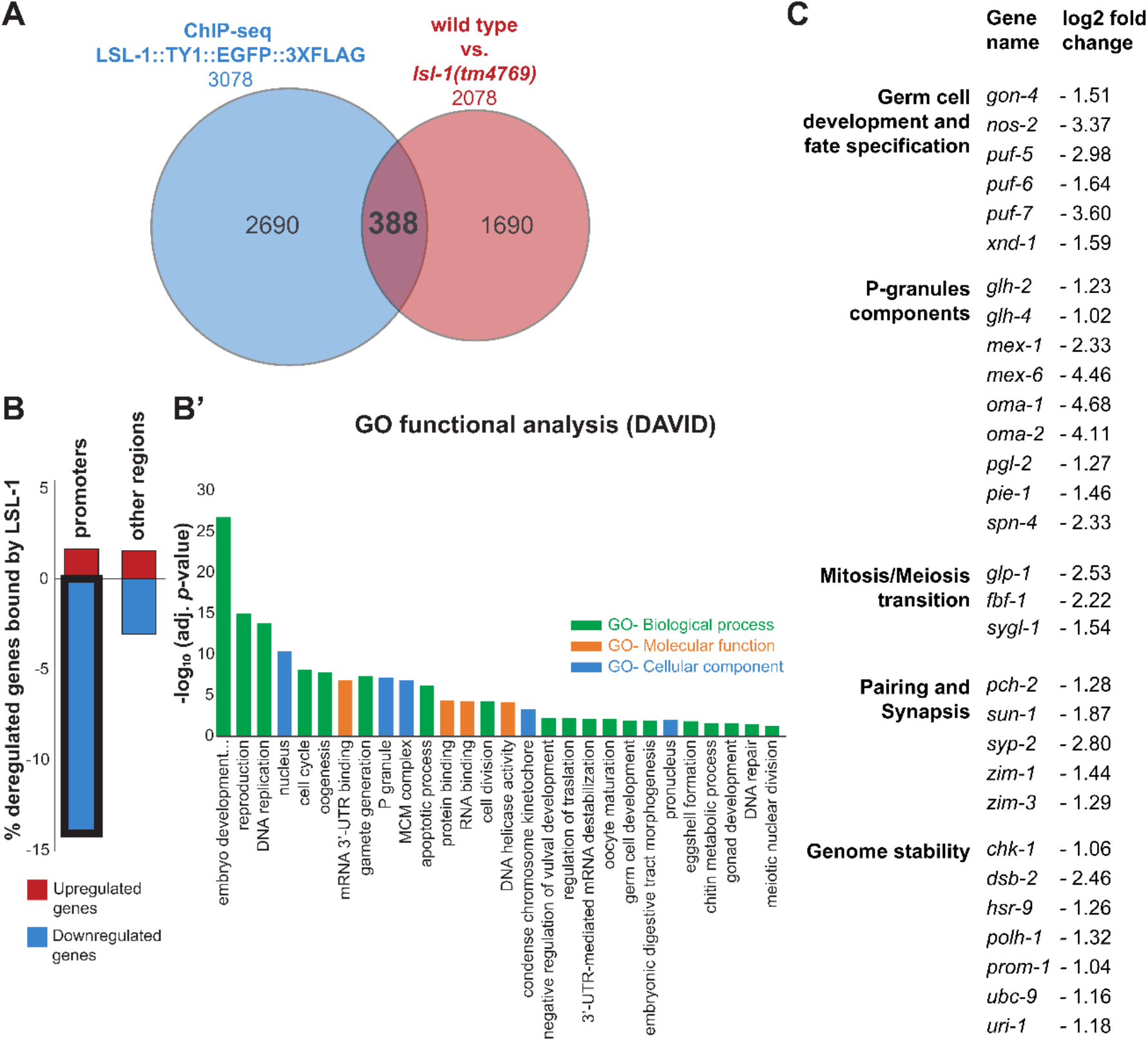
LSL-1 acts mainly as a transcriptional activator of germline genes. (A) Overlap between LSL-1::TY1::EGFP::3xFLAG Chip-seq data from modERN resource (Kudron *et al*. 2018) and RNA-seq analysis data of *lsl-1 (tm4769)* DEG with respect to wild type. Common intercepts are significant DEGs (*q*-value ≤ 0.01, −2 ≥ FC ≥ 2) and significant LSL-1::TY1::EGFP::3xFLAG binding sites (IDR ≤ 0.1%). Overlap was significant (*p*-value ≤ 0.0001). Statistical significance was assessed using cross comparation contingency tables by chi-square test with Yates correction (see Table S2). (B) Graph illustrates the percentage of the 388 significant DEG in *lsl-1 (tm4769)* animals with respect to wild type and, simultaneously, LSL-1::TY1::EGFP::3xFLAG target genes, distributed by LSL-1 binding site (promoter vs. other regions). Thick line depicts that most significant DEGs are downregulated and directly bound by LSL-1 in their promoter region. (B’) Histogram shows DAVID GO term functional analysis (Huang *et al*. 2009) for downregulated genes in *lsl-1 (tm4769)* animals with respect to wild type and bound by LSL-1 in their promoter region. Data are plotted as vertical bars that represent the significance of each GO-term. Adj. *p*-values were all significant (adj. *p*-value ≤ 0.05) and correspond to Benjamini–Hochberg correction. (C) Illustration shows representative germline genes downregulated in *lsl-1(tm4769)* animals with respect to wild type and bound by LSL-1 in their promoter region, clustered by their associated germline function. Note, symbol (-) in fold change column stands for downregulation in the RNA-seq analysis. DEGs, differentially expressed genes; GO, gene ontology.

Data presented herein indicate that LSL-1 could function as a direct transcriptional activator of germline genes involved in different processes, ranging from germ cell maintenance to pairing and synapsis processes, DNA stability, and P-granules composition (Figure 7C).

### LSL-1 functions by antagonizing LET-418/Mi2 and HPL-2 /HP1 heterochromatin proteins

*lsl-1* was identified as a suppressor of developmental defects associated with mutations in *let-418*, which encodes a chromatin remodeler (Erdelyi *et al*. 2017). To test whether *lsl-1* and *let-418* were also genetically interacting in the germline, we generated *lsl-1; let-418* double mutants and measured their fertility. At the restrictive temperature of 22 °C, fertility and embryo viability are partially restored in *let-418(n3536)* double mutant combination with the *lsl-1(ljm1)* allele, indicating that LSL-1 might antagonize LET-418/Mi2 function in the germline. LET-418/Mi2 is part of chromatin proteins known to repress transcription (Ahringer and Gasser 2018). To further investigate whether LSL-1 functions by antagonizing repressive chromatin formation, we investigated the genetic interaction of *lsl-1* with heterochromatin factor coding genes *hpl-1* and *hpl-2*, and histone H3K9 methyltransferase coding genes *met-2* and *set-25* (Ahringer and Gasser 2018). The H3K9 methyltransferases SET-25 and MET-2 are known to be responsible for most genomic H3K9 methylation (Towbin *et al*. 2012). We generated *lsl-1; met-2 set-25* triple mutants; however, loss of H3K9 methylation does not rescue *lsl-1* mutant phenotype, nor does absence of HPL-1 activity restore fertility. Nevertheless, lack of HPL-2/HP1 activity partially restores fertility and confers viability to the embryos (Table 3). These overall genetic interactions suggest that LSL-1 could activate germline gene transcription by antagonizing repressive chromatin formation by LET-418/Mi2 and HPL-2/HP1.

**Table 3.** Genetic interactions of *lsl-1* with chromatin factor genes.

| Genotype | Mean brood size<sup>a</sup> | Viability (%) | Incidence of males (%) | *n*<sup>b</sup> |
| --- | --- | --- | --- | --- |
| wild type | 301.79 ± 31.25 | 98.44 | 0.06 | 43 |
| *lsl-1(tm4769)* | 28.40 ± 23.95 | 0.12 | n/a | 59 |
| *lsl-1(ljm1)* | 58.26 ± 27.18 | 5.41 | 19.31 | 46 |
| *hpl-1(tm1624)* | 287.13 ± 31.73 | 94.43 | 0.05 | 16 |
| *lsl-1(tm4769); hpl-1(tm1624)* | 45.50 ± 29.07<sup>n.s</sup> | 0.00 | n/a | 12 |
| *lsl-1(ljm1); hpl-1(tm1624)* | 58.58 ± 42.29<sup>n.s</sup> | 2.42 | 41.18 | 12 |
| *hpl-2(tm1489)* | 248.85 ± 20.26 | 97.29 | 0.00 | 20 |
| *lsl-1(tm4769); hpl-2(tm1489)* | 128.82 ± 41.62<sup>***</sup> | 14.82 | 6.19 | 11 |
| *lsl-1(ljm1); hpl-2(tm1489)* | 153.64 ± 53.32<sup>***</sup> | 39.64 | 5.67 | 11 |
| *met-2(n4256) set-25(n5021)* | 276.62 ± 58.12 | 98.62 | 0.00 | 21 |
| *lsl-1(tm4769); met-2(n4256) set-25(n5021)* | 1.33 ± 3.47<sup>***c</sup> | 0.00 | n/a | 12 |
| *lsl-1(ljm1); met-2(n4256) set-25(n5021)* | 53.73 ± 26.65<sup>n.s</sup> | 6.94 | 9.76 | 11 |
| wild type<sup>22 °C</sup> | 241.33 ± 46.45 | 98.62 | 0.07 | 6 |
| *lsl-1(tm4769)*<sup>22 °C</sup> | 20.25 ± 19.71 | 0.00 | n/a | 12 |
| *lsl-1(ljm1)*<sup>22 °C</sup> | 8.75 ± 20.42 | 10.48 | 9.09 | 12 |
| *let-418(n3536)*<sup>22 °C</sup> | 147.33 ± 67.25 | 95.25 | 0.23 | 12 |
| *lsl-1(tm4769); let-418(n3536)*<sup>22 °C</sup> | 0.58 ± 1.16<sup>**c</sup> | 0.00 | n/a | 12 |
| *lsl-1(ljm1); let-418(n3536)*<sup>22 °C</sup> | 17.31 ± 16.80 | 32.44 | 4.69 | 13 |

<sup>a</sup> Data correspond to the mean ± SD of the total number of eggs laid per hermaphrodite parent. Statistical comparison between *lsl-1* double or triple mutants and their corresponding *lsl-1* genotype at the defined temperature, performed by two-tailed Student's *t*-test with Welch's correction. \*\*\* *p*-value ≤ 0.001, \*\* *p*-value ≤ 0.01, <sup>n.s</sup> *p*-value > 0.05.
<sup>b</sup> Total number of parental hermaphrodites per genotype.
<sup>c</sup> Statistical significance does not indicate *lsl-1* fertility rescue.
n/a not applicable

## DISCUSSION

This study defines the *C. elegans* protein LSL-1 as a novel crucial transcriptional activator of germline genes. LSL-1 is present at all stages of germline development, from primordial germ cells proliferation to differentiation through meiotic prophase progression. *lsl-1* loss-of-function mutants produce almost no functional gametes as a result of chromosome pairing defects, defective meiotic recombination, and genome instability. Transcriptomic analysis and ChIP-seq data show that LSL-1 binds germline gene promoters, acting mainly as a transcriptional activator. Furthermore, our genetic interaction analyses reveal that LSL-1 antagonizes the function of the heterochromatin proteins HPL-2/HP1 and LET-418/Mi2 to ensure production of viable progeny.

*lsl-1* encodes a C2H2-type zinc-finger transcription factor closely homologue to LSY-2. *lsy-2* is expressed ubiquitously and is required for the specification of ASE neurons, for proper vulva patterning, and for repression of germline genes in somatic cells (Johnston 2005; Lin *et al*. 2015). LSL-1, in contrast, appears to be expressed specifically in the germline. Both proteins are members of the SP1/KLF family of transcription factors, involved in growth and developmental processes (Figure S1) (Suske *et al*. 2005; Kim *et al*. 2017). These share a triple C2H2-type zinc-finger cluster and a less conserved double zinc finger at the C-terminus. The closest human homolog is ZFP57, which contains an additional KRAB domain that interacts with the heterochromatin protein 1 (HP1) (Figure S1) (Li *et al*. 2008; Quenneville *et al*. 2011). Protein sequence and structure analysis suggest that LSL-1 is a DNA binding protein whose function is not conserved.

In the present study, two different alleles were used: *tm4769* and *ljm1* (Figure 1A). Although worms bearing one or the other allele exhibit very similar defects, the penetrance of the phenotype is different despite repeated backcrosses performed to eliminate any additional mutations. A possible interpretation is that a cryptic translational initiation site (TIS) is used by the ribosome in *lsl-1(ljm1)* mutants to produce a protein that still retains some functionality. Two AUG codons downstream of the predicted TIS exhibit conserved nucleotides at position −3, −2, and +4 that could function as TIS consensus sequences (Hernández *et al*. 2019). Detailed comparison of the transcriptome of both mutant alleles (see below) is consistent with the interpretation that *ljm1* represents a hypomorphic allele. Additional deregulated genes observed in *lsl-1(tm4769)* with respect to *lsl-1(ljm1)* are also enriched in germline genes, and most of them are bound by LSL-1 in their promoter region (File S1).

LSL-1 is first detected in the P4 blastomere and could potentially represent the initial transcription factor that activates zygotic transcription of germline genes. The first zygotic germline transcripts, including LSL-1 targets, have been detected in Z2 and Z3 PGCs (Wang and Seydoux 2013; Lee *et al*. 2017). An interesting hypothesis would be that LSL-1 is part of the process that initiates the germline transcriptional program by interpreting the epigenetic memory of germline transcription. Germline genes are marked in the parental germline by the histone methyltransferase MES-4, which deposits H3K36 methyl marks while germline genes are transcribed (Tursun *et al*. 2011; Patel *et al*. 2012). These marks are transmitted and maintained in the embryos by MES-4 maternal contribution and therefore constitute an epigenetic memory of germline transcription. However, LSL-1 would function redundantly with other factors since its absence still leads to germ cell proliferation and a certain degree of germ cell differentiation.

LSL-1 appears to be one of the few transcriptional regulators functioning in the germline, LAG-1/CSL being another key transcription factor that controls germ cell fate in response to Notch signaling (Chen *et al*. 2020). To date, crucial studies have shown that proper patterning of germline gene expression requires the 3’UTR (Merritt *et al*. 2008). LSL-1 could function as a general activator of germline genes transcription followed by fine-tuning at the post-transcriptional level for proper patterning.

Monitoring of chromosome pairing by FISH or HIM-8, and SUN-1 localization showed that pairing dynamics are impaired in *lsl-1* mutants (Penkner *et al*. 2007; Woglar *et al*. 2013). Recombination also appears defective, as revealed by a high number of univalent in oocyte nuclei and an increased level of apoptotic germ cells, which depends on the recombination initiator protein SPO-11 (Dernburg *et al*. 1998). These observations are in agreement with our transcriptomic analysis. A large number of genes encoding essential meiotic proteins are downregulated in the absence of LSL-1, including the chromosomal axis component HTP-1, the synaptonemal complex proteins SYP-2 and SYP-4 (Martinez-Perez 2005), or the pairing center binding ZIM proteins (ZIM-1, −2, −3, and HIM-8) (Phillips and Dernburg 2006). Genes involved in recombination, DNA repair, and genome stability, such as *dsb-2*, which plays a role in the control of meiotic DSB formation, or *hsr-9* and *chk-1*, which are involved in the cell cycle checkpoints regulation in response to DNA damage, were also found to be downregulated in *lsl-1* mutants (Ryu *et al*. 2013; Rosu *et al*. 2013; Zhang and Hunter 2014). These observations together with the germline expression pattern throughout development indicate a general role of LSL-1 in the transcription regulation of germline genes. However, all meiotic processes described above, including pairing, synapsis, and recombination, are significantly compromised but not completely abolished in the absence of LSL-1. A possible interpretation is that LSL-1 acts redundantly with other regulators and, in the absence of LSL-1, the stoichiometry of key factors involved in pairing, recombination and genome stability might be highly perturbed.

Defects associated with mutations in *lsl-1* are mediated partially by the *C. elegans* HPL-2/HP1 heterochromatin protein and the chromatin remodeler LET-418/Mi2 (von Zelewsky *et al*. 2000; Bannister *et al*. 2001; Dialynas *et al*. 2008). LSL-1 was identified in a screen for suppressors of developmental defects associated with the absence of the chromatin repressor LET-418/Mi2. LSL-1 is required for ectopic localization of P granules in somatic cells of *let-418* mutants (Erdelyi *et al*. 2017). The present study reveals that LSL-1 also antagonizes LET-418/Mi2 function in the germline. Partial fertility is restored (Table 3), and we could also observe a significant decrease in the length of the transition zone in *lsl-1; let-418* double mutants (data not shown). The heterochromatin protein HPL-2/HP1 also contributes to *lsl-1* phenotype. A global reorganization of chromatin could take place in the *lsl-1* mutant germline, where HPL-2/HP1 and LET-418/Mi2 play major roles. Similarly to LET-418/Mi2, HPL-2/HP1 is known to act as a repressor of germline gene expression in the somatic cells (Coustham *et al*. 2006; Meister *et al*. 2011). This interaction of *lsl-1* with *hpl-2* and *let-418* suggests that, in the absence of LSL-1, the germline chromatin adopts a conformation resembling the somatic one. However, no large set of somatic genes are upregulated in *lsl-1* mutants (File S2), indicating that downregulation of germline genes is not accompanied by somatic gene expression, at least not at the stage examined.

In conclusion, we characterize herein a new transcriptional regulator of genes that are involved in a wide range of germline processes. Since *lsl-1* is starts in the P4 blastomere, we propose that LSL-1 might initiate the germline transcriptional program and might be part of the process that interprets the epigenetic memory established in the parental germline by antagonizing HPL-2/HP1 and LET-418/Mi2 function, specifically in the germ cells. Identifying the mechanisms by which LSL-1 is recruited to the chromatin will contribute to understand how transcriptional programs are triggered in development and disease.

## Supporting information

List of supplemental materials

## DATA AVAILABILITY STATEMENT

Sequencing files of the LSL-1 ChIP-seq experiment are accessible through the modERN website http://epic.gs.washington.edu/modERN/. Raw sequencing files of the RNA-seq experiment have been deposited in the ArrayExpress database at EMBL-EBI (www.ebi.ac.uk/arrayexpress) under accession number E-MTAB-11199. Strains and reagents used in this study are available upon request. Supplemental Materials consisting of supplemental methods, supplemental figures, supplemental tables and supplemental files have been deposited at figshare portal https://gsajournals.figshare.com/submit.

## ACKNOWLEDGEMENTS

The authors thank L. Schild and L. Buillard for excellent technical support; the “model organism Encyclopedia of Regulatory Networks” resource (modERN; http://epic.gs.washington.edu/modERN/) for ChIP-seq data; A.F. Dernburg for providing the guinea pig α-HIM-8 antiserum; M. Zetka for providing the rabbit α-HTP-3 antibody; M. Boxem for providing the guide RNA cloned from the plasmid pMB70.

Some strains were provided by the *Caenorhabditis* Genetics Center (CGC; cbs.umn.edu/cgc/home), which is funded by NIH Office of Research Infrastructure Programs (P40 OD010440), and the National BioResource Project (NBRP; http://www.shigen.nig.ac.jp/c.elegans), *C. elegans* Gene Knockout Consortium by S. Mitani at the Tokyo Women’s Medical University School of Medicine (Tokyo, Japan). Some strains used in this study were provided by A.M. Villeneuve at Stanford University School of Medicine (Stanford CA, USA) and J. Ceron at *C. elegans* Core Facility-IDIBELL (Barcelona, Spain).

