## Supplementary material for "The zinc-finger transcription factor LSL-1 is a major regulator of the germline transcriptional program in *C. elegans*": List of supplemental materials

### Supplemental Methods

List of strains used in this study

LSL-1 domains prediction and protein alignments

List of antibodies used in this study

Immunofluorescence in adult hermaphrodite gonads

Germline mitotic region cytological analysis

Fluorescence in situ hybridization (FISH)

Acridine orange (AO) staining

RNA-seq data analysis

Chromatin immunoprecipitation (ChIP), data processing, and analysis

Functional analysis

Supplemental Methods Bibliography

### Supplemental Figures

Figure S1 *lsl-1* encodes a 318 aa Zinc-finger C2H2-type protein

Figure S2 LSL-1 homologous proteins

Figure S3 *lsl-1(ljm1)* worms exhibit *lsl-1(tm4769)* comparable altered chromatin organization in the germline and abnormal progression through meiotic prophase

Figure S4 LSL-1 is required for the proper progression of homologous chromosome pairing

Figure S5 Analysis of the mitotic region

Figure S6 Apoptosis in *lsl-1(ljm1)* mutants

Figure S7 Germline genes expression changes in *lsl-1(ljm1)* mutants

Figure S8 Transgene *wgIs720[lsl-1::TY1::EGFP::3xFLAG]* resembles the *lsl-1* expression pattern

Figure S9 Motif analysis of LSL-1::TY1::EGFP::3XFLAG significant ChIP-seq peaks

Figure S10 LSL-1 acts mainly as a transcriptional activator of germline genes

Supplemental Figures Bibliography

**Supplemental Figures**

Figures S1 to S10

**Supplemental Tables**

Table S1 Brood size, survival rate, and incidence of males determination for both *lsl-1* alleles (at 22 °C and 25 °C)

Table S2 Cross-comparison contingency tables summary

### Supplemental Files

File S1 DEGs summary

File S2 TEA summary

File S3 DEGs in *ljm1* and *tm4769* alleles direction

File S4 LSL-1 ChIPseq results

File S5 Overlap DEGs in *lsl-1* mutants and LSL-1 bound genes
