## Supplemental figures for "The zinc-finger transcription factor LSL-1 is a major regulator of the germline transcriptional program in *C. elegans*"

This PDF file includes:

Figures S1 to S10

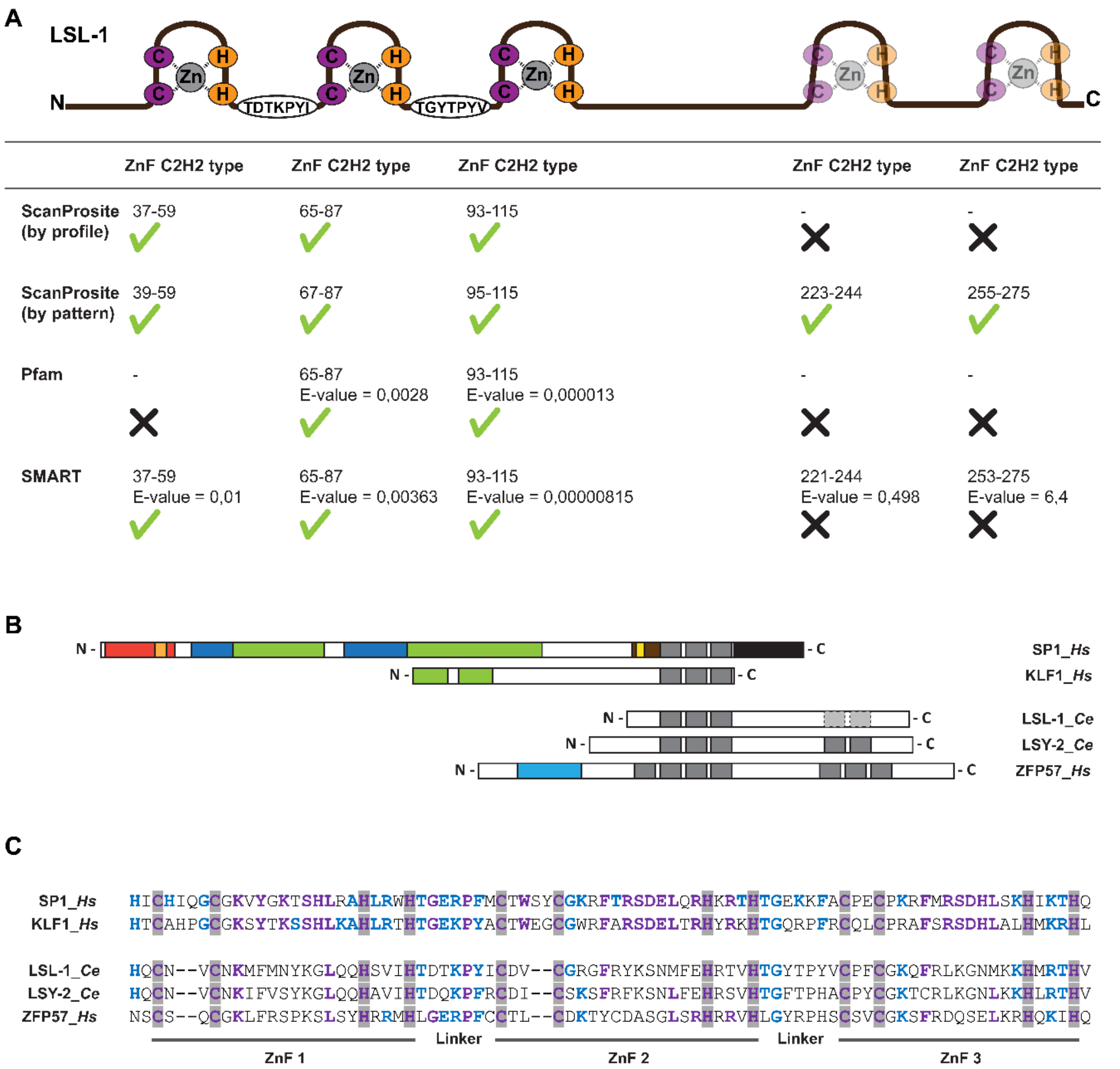
A
LSL-1
N
C
Zn
TDTKPYI
TGYTPYV
ZnF C2H2 type
ScanProsite (by profile)
37-59
65-87
93-115
ScanProsite (by pattern)
39-59
67-87
95-115
223-244
255-275
Pfam
65-87 E-value = 0,0028
93-115 E-value = 0,000013
SMART
37-59 E-value = 0,01
65-87 E-value = 0,00363
93-115 E-value = 0,00000815
221-244 E-value = 0,498
253-275 E-value = 6,4
B
SP1_Hs
KLF1_Hs
LSL-1_Ce
LSY-2_Ce
ZFP57_Hs
C
SP1_Hs HICHIQGCGKVYGKTSHLRAHLRWHTGERPFMCTWSYCGKRFTRSDELQRHKRTHTGEKKFACPECPKRFMRSDHLSKHIKTHQ
KLF1_Hs HTCAHPGCGKSYTKSSHLKAHLRTHTGEKPYACTWEGCGWRFARSDELTRHYRKHTGQRPFRCQLCPRAFSRSDHLALHMKRHL
LSL-1_Ce HQCN--VCNKMFMNYKGLQQHSVIHTDTKPYICDV--CGRGFRYKSNMFEHRTVHTGYTPYVCPFCGKQFRLKGNMKKHMRTHV
LSY-2_Ce HQCN--VCNKIFVSYKGLQQHAVIHTDQKPFRCDI--CSKSFRFKSNLFEHRSVHTGFTPHACPYCGKTCRLKGNLKKHLRTHV
ZFP57_Hs NSCS--QCGKLFRSPKSLSYHRRMHLGERPFCCTL--CDKTYCDASGLSRHRRVHLGYRPHSCSVCGKSFRDQSELKRHQKIHQ
ZnF 1
Linker
ZnF 2
Linker
ZnF 3

**Figure S1** *lsl-1* encodes a 318 aa Zinc-finger C2H2-type protein. (A) Schematic representation of the LSL-1 protein. Three first predicted Zinc-fingers (ZnF) C2H2-type show conserved cysteines in violet and histidines in orange chelating a single Zinc ion in grey. The two C-terminal ZnF (only predicted by the ScanProsite by pattern resource) are indicated in light colors. Table presents the domains predicted for LSL-1 full-length sequence by the indicated bioinformatic tools. Note, (✓) symbol stands for significant predicted ZnF C2H2-type domain, while (✖) symbol stands for non-predicted ZnF C2H2-type domain. (B) Graph illustrates the comparison between the primary structures of human SP/KLF protein family members (SP1 and KLF1) and the LSL-1 protein and LSL-1 homologous proteins (LSY-2 and ZFP57). Both SP/KLF proteins contain the highly conserved cluster of three ZnF C2H2-type that includes the DNA-binding domain and is shared by LSL-1 and LSL-1 homologs (grey boxes). Illustration shows, in addition, repressor domains (red boxes), transactivation (Q-rich) domains (green boxes), S/T rich domains (dark-blue boxes), and the conserved SP sequence motifs Sp box (orange) and BTD (yellow box). KRAB DNA-binding domain is indicated as a light-blue box (O'Connor *et al.* 2016; Perkins *et al.* 2016). (C) Alignment of the first three ZnF and linkers of LSL-1 with SP/KLF family representative members SP1 and KLF1, and LSL-1 homolog proteins LSY-2 and ZFP57. Conserved cysteines and histidines of the three ZnF shown in grey shadow. Amino acids in violet bold letters are 100% conserved, while aa in blue bold letters are partly conserved (> 70%). ZnF, Zinc finger; Ce, *C. elegans*; Hs, *Homo sapiens.*

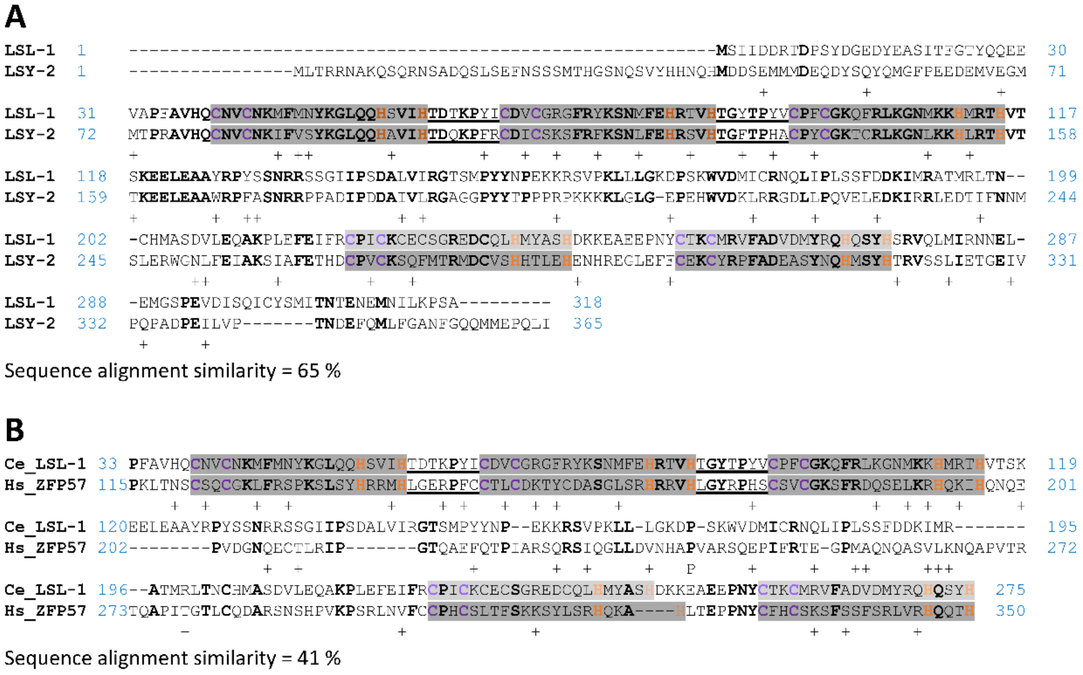


**Figure S2** LSL-1 homologous proteins. (A and B) Sequence alignment of LSL-1 and (A) its paralog LSY-2 and (B) its human homolog ZPF57. Zinc-fingers C2H2-type indicated in grey boxes, with conserved cysteines in violet and histidines in orange. Identical aa are indicated in bold letters. Ce, *C. elegans*; Hs, *Homo sapiens*; +, aa with similar function.

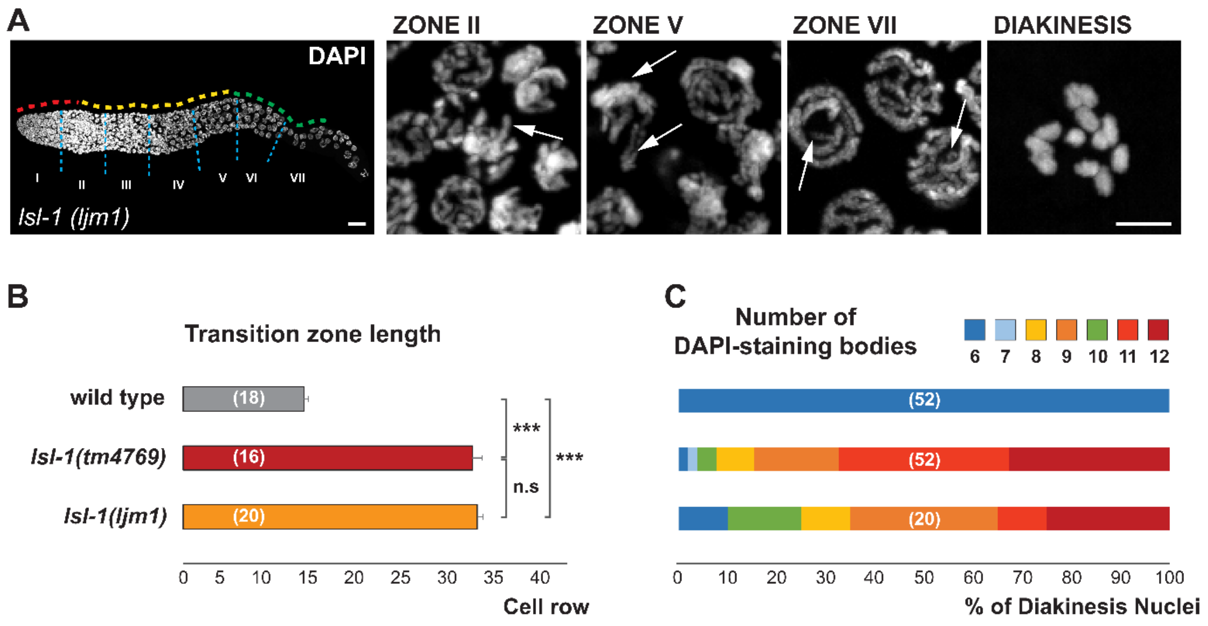


Figure S3 *lsl-1(ljm1)* worms exhibit *lsl-1(tm4769)* similar altered chromatin organization in the germline and abnormal progression through meiotic prophase. (A) Representative confocal projection images of DAPI-stained (grey) gonads from dissected *lsl-1(ljm1)* 1-day-old adult hermaphrodites. Each panel shows a magnification of the indicated zones. Dashed lines depict the mitotic region (red), the transition zone to meiosis (yellow), the pachytene stage (green), and the boundaries of the seven equally long zones (light blue). Arrows point to altered chromatin structures (see text). (B) Graphic representation of the transition zone length, determined by DAPI staining and quantified in nuclei rows from the MR/TZ boundary to the TZ/PS limit. Data are plotted as horizontal bars that represent mean length. Error bars correspond to standard error (SEM). *p*-value ≤ 0.001 (***); *p*-value > 0.05 nonsignificant (n.s), by two-tailed Student's *t*-test with Welch's correction. Number of germlines scored for each genotype in brackets. (C) Percentages of diakinetic oocytes by number of DAPI-staining bodies content in 1-day adult hermaphrodite germlines for the indicated genotypes. Note, wild type and *lsl-1(tm4769)* data added to the graphics for clearer *lsl-1(ljm1)* results interpretation. Number of oocytes scored for each genotype in brackets. Scalebars, 20 µm and 5 µm in whole gonad images and magnification panels, respectively. MR, mitotic region; TZ, transition zone; PS, pachytene stage.

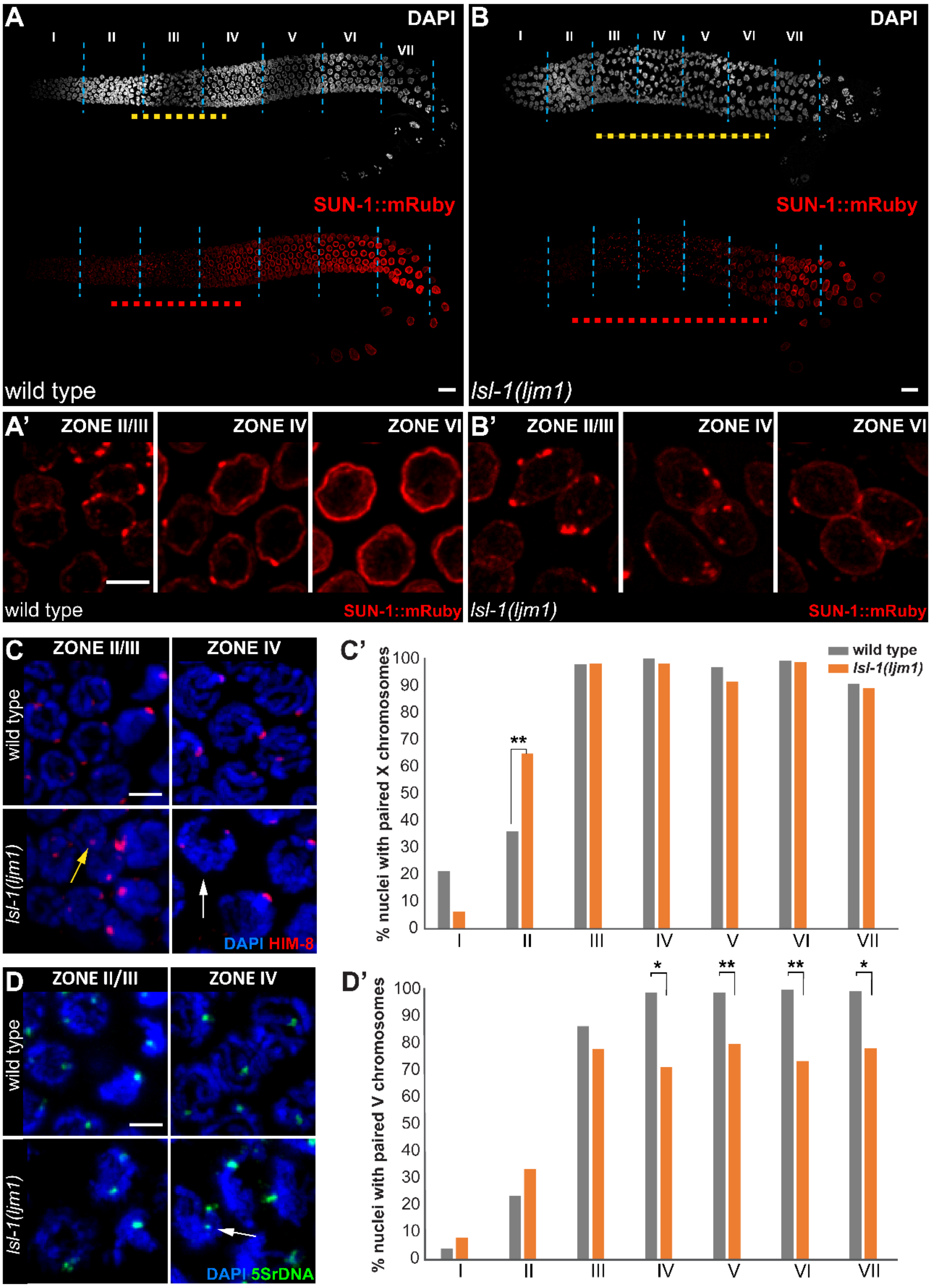
A
DAPI
I
II
III
IV
V
VI
VII
SUN-1::mRuby
wild type
B
DAPI
SUN-1::mRuby
lsl-1(ljm1)
A'
ZONE II/III
ZONE IV
ZONE VI
wild type
SUN-1::mRuby
B'
ZONE II/III
ZONE IV
ZONE VI
lsl-1(ljm1)
SUN-1::mRuby
C
ZONE II/III
ZONE IV
wild type
lsl-1(ljm1)
DAPI HIM-8
C'
% nuclei with paired X chromosomes
wild type
lsl-1(ljm1)
**
0
10
20
30
40
50
60
70
80
90
100
I
II
III
IV
V
VI
VII
D
ZONE II/III
ZONE IV
wild type
lsl-1(ljm1)
DAPI 5SrDNA
D'
% nuclei with paired V chromosomes
*
**
**
*

**Figure S4** LSL-1 is required for the proper progression of homologous chromosome pairing. (A and B) Representative confocal projection images of 1-day-old adult stage gonads of (A) wild-type and (B) *lsl-1(ljm-1)* animals, expressing SUN-1::mRuby (red) and stained with DAPI (grey). Dashed lines delineate nuclei showing SUN-1::mRuby patches (red), extension of the transition zone (yellow), and the boundaries between the seven equally long zones (light blue). (A' and B') Each panel represents a magnification of the indicated zones. (C and D) Representative images of zone II/III and zone IV nuclei of the indicated genotypes: (C) immunostained with HIM-8 antibody (red); (D) hybridized with 5S rDNA FISH probe (green) to monitor chromosome pairing and costained with DAPI (blue). Arrows point to possible precocious paired chromosomes in late mitotic zone (yellow) or nuclei with unpaired signals (white). (C' and D') Histograms showing the percentage of nuclei with paired (C') HIM-8 and (D') 5S rDNA signals, scored per zones of the indicated genotypes. $p$-value ≤ 0.001 (***); $p$-value ≤ 0.01 (**); $p$-value ≤ 0.05 (*); $p$-value > 0.05 nonsignificant, by two-tailed Student's $t$-test with Welch's correction. At least three gonads from independent experiments were scored for each genotype. Scale bars, 20 µm and 5 µm in whole gonad images and magnification panels, respectively.

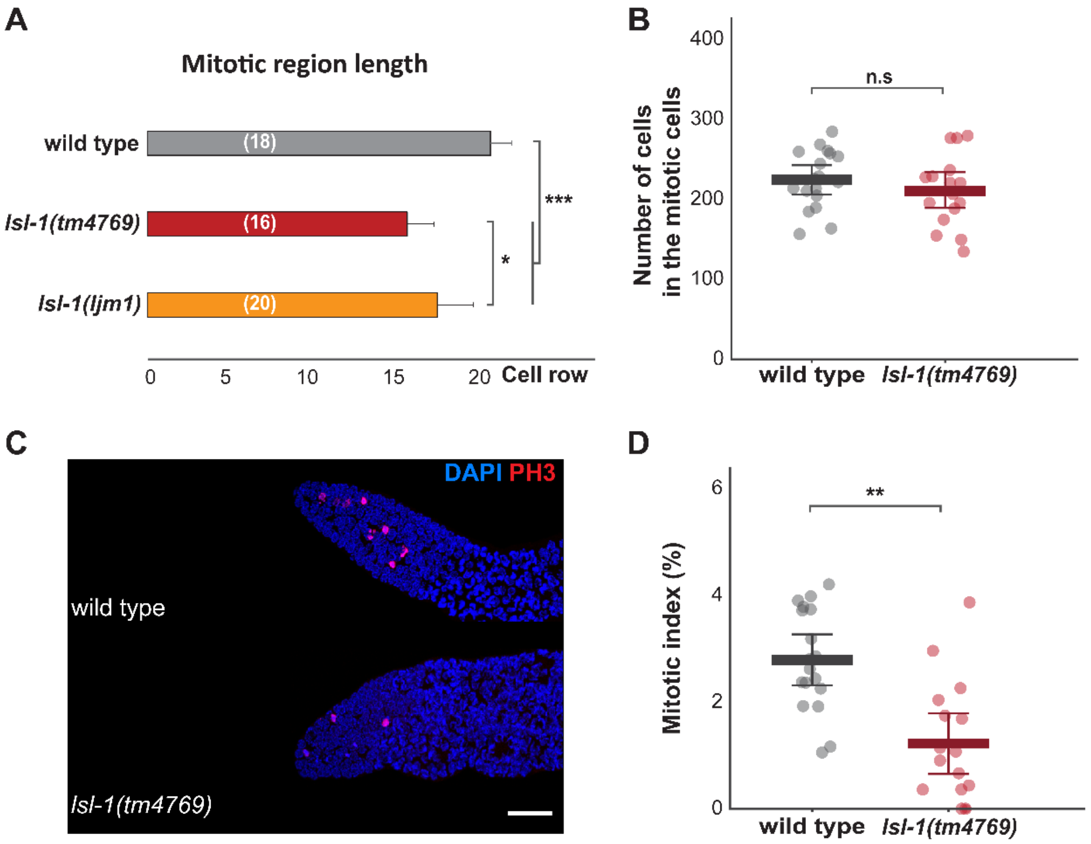
A
Mitotic region length
wild type
lsl-1(tm4769)
lsl-1(ljm1)
(18)
(16)
(20)
***
*
0
5
10
15
20
Cell row
B
Number of cells in the mitotic cells
400
300
200
100
0
n.s
wild type
lsl-1(tm4769)
C
DAPI PH3
wild type
lsl-1(tm4769)
D
Mitotic index (%)
6
4
2
0
**
wild type
lsl-1(tm4769)

**Figure S5** Analysis of the mitotic region. (A) Graphic representation of the length extension of the mitotic region quantified in nuclei rows from the distal tip cell to the TZ limit, from 1-day-old adult hermaphrodite gonads for wild-type and *lsl-1* genotypes. The length of the mitotic region was determined based on chromatin morphology assessed by DAPI staining. Data are plotted as horizontal bars that represent mean length. Error bars correspond to standard error (SEM). *p*-value ≤ 0.001 (***); *p*-value ≤ 0.05 (*), by two-tailed Student's *t*-test with Welch's correction. Number of germlines scored for each genotype in brackets. (B) Scatter plot illustrating the total number of nuclei in the mitotic region of the same specimens as in (A) for wild-type and *lsl-1(tm4769)* genotypes. Data are plotted as vertical dot plots, with each dot representing the number of nuclei in the mitotic region of one gonad. Horizontal lines represent mean, with error bars corresponding to standard deviation (SD). *p*-value > 0.05 nonsignificant (n.s), by two-tailed Student's *t*-test with Welch's correction. (C) Representative confocal projection images of the *C. elegans* mitotic region from 1-day-old adult hermaphrodite gonads for the indicated genotypes, immunostained using antibodies against α-Histone H3 Ser10-p (PH3) to label mitotic nuclei (red) and costained with DAPI (blue). (D) Scatter plot showing the mitotic index estimated in the same specimens as in (A) for the indicated genotypes. Note, mitotic index is defined here as the number of PH3 positive cells over the total number of nuclei in the mitotic region and represented as a percentage. Data are plotted as vertical dot plots, with each dot representing mitotic index value of one gonad. Horizontal lines represent mean with error bars corresponding to standard deviation (SD). *p*-value ≤ 0.01**, by two-tailed Student's *t*-test with Welch's correction. Scalebar, 20 µm.

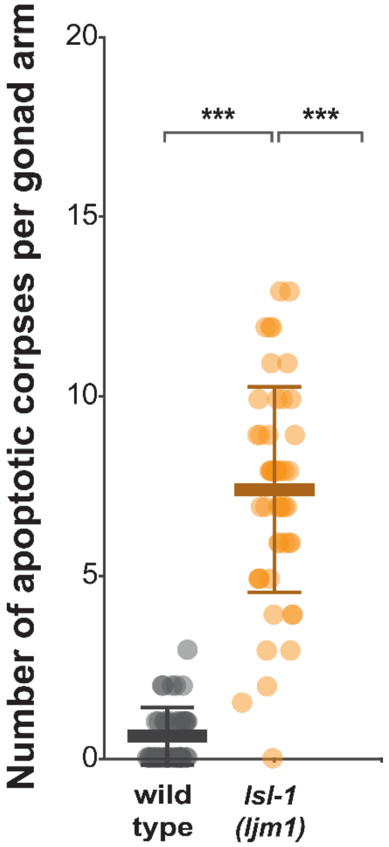


**Figure S6** Apoptosis in *lsl-1(ljm1)* mutants. Scatter plot showing the number of apoptotic corpses per gonad arm from 1-day adult hermaphrodites for the indicated genotypes and quantified by acridine orange staining. Data are plotted as vertical dot plots, with each dot representing the number of apoptotic corpses in one gonad arm. Horizontal lines represent mean, with error bars corresponding to standard deviation (SD). *p*-value ≤ 0.001***; *p*-value > 0.05 nonsignificant (n.s), by two-tailed Student's *t*-test with Welch's correction. At least 24 gonads from different biological replicates were scored for each genotype.

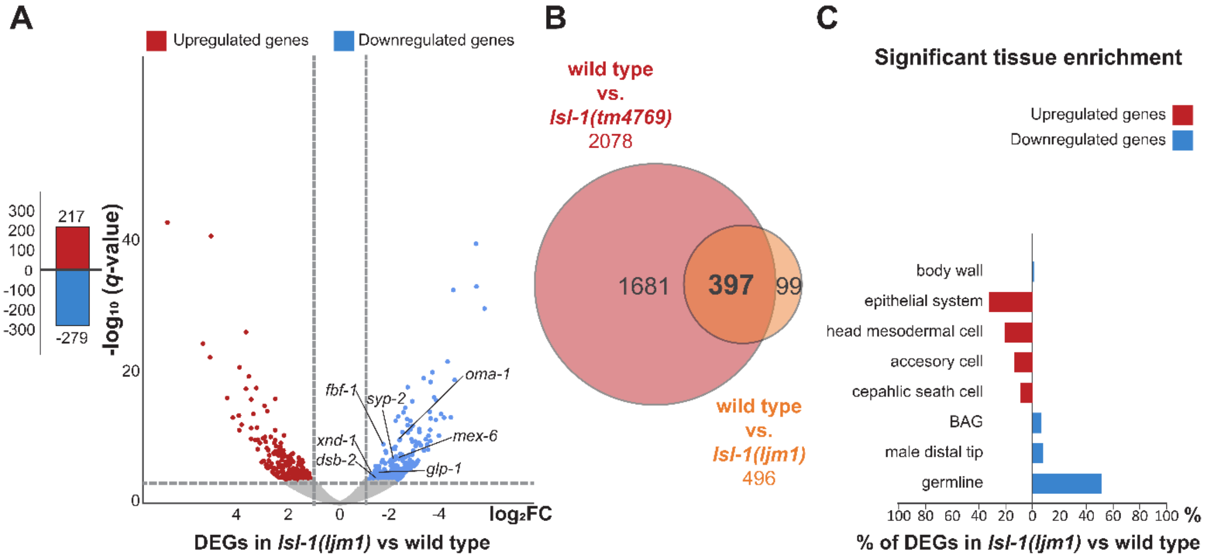


**Figure S7** Germline genes expression changes in *lsl-1(ljm1)* mutants. (A) Bar and volcano plots show the number of significant differentially expressed genes (DEGs) in *lsl-1(ljm1)* young adult animals compared with wild type, determined by RNA-seq analysis. Each dot represents a gene, and red and blue colors correspond to significant up- and downregulated genes, respectively. Dash lines indicate the significance and fold change cutoffs ($q$-value $\leq$ 0.01 and $-2 \geq$ fold change $\geq 2$). In italics, representative genes associated to different germline functions (see results section). Note, symbol (-) stands for downregulation both in number of genes and fold change; $q$-value stands for adjusted $p$-value found using an optimized FDR approach (Storey and Tibshirani 2003). (B) Venn diagram illustrating the cross comparison between number of DEGs in *lsl-1(tm4769)* young adult animals compared with wild type (red), and the number of DEGs in *lsl-1 (ljm1)* young adult animals compared with wild type (orange). Statistical significance for the 397 overlapped genes was assessed using cross comparison contingency tables by chi-square test with Yates correction ($p$-value $\leq 0.0001$). (C) Graph illustrating the tissue enrichment of significant up- (red) and downregulated (blue) genes in *lsl-1(ljm1)* young adult animals compared with wild type, using the T.E.A-Wormbase tool (Angeles-Albores *et al.* 2016) and represented as a percentage of total significant up- or downregulated genes. Only enrichments with significant adj. $p$-value $\leq 0.05$ were scored. For enrichment in blastomeres, see File S2. FC, fold change; DEGs, differentially expressed genes; T.E.A, tissue enrichment analysis; BAG, neuron class of two neurons with ciliated endings, in the head, with elliptical, closed, sheet-like processes near the cilium, which envelop a piece of hypodermis (see Wormbase anatomy term).

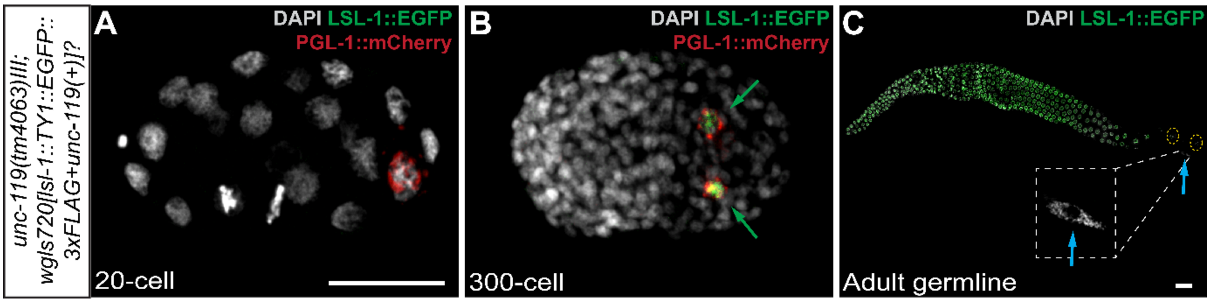


**Figure S8** Transgene *wgIs720[lsl-1::TY1::EGFP::3xFLAG]* expression matches the *lsl-1* expression pattern (Figure 1). (A–C) Representative confocal projection images of *C. elegans* at different developmental stages. Chromatin is stained with DAPI (grey), PGL-1::mCherry (red) marks germ cells, and LSL-1::EGFP or LSL-1::GFP (green) reflects the endogenous *lsl-1* expression. Green arrows point to precursor cells Z2/Z3, and light-blue arrows, to the somatic cells of the adult gonad. (A) LSL-1 was not detected in embryos before the 24-cell stage. (B) *lsl-1* is expressed in the PGC during embryonic development and maintained in germ cells in (C) adulthood. Scalebars, 20 µm.

| Motif | Logo | Reverse Complement Logo | E-value |
|---|---|---|---|
| TACBGTA | 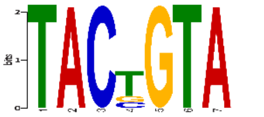 | 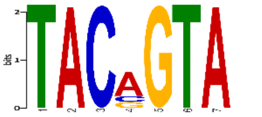 | 3.1e-369 |
| GGTCTCR | 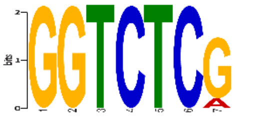 | 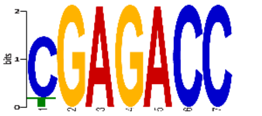 | 1.8e-122 |
| AAASGCGC | 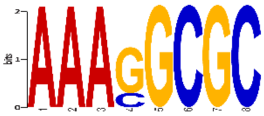 | 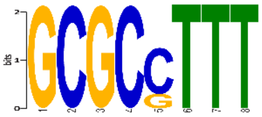 | 3.6e-121 |
| GCANACAC | 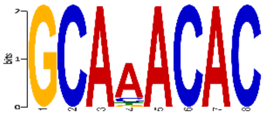 | 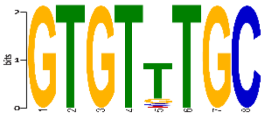 | 1.9e-73 |
| YBHDYM | 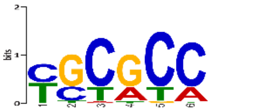 | 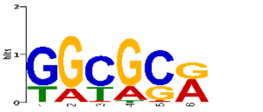 | 9.4e-55 |
| ATWTTY | 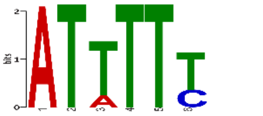 | 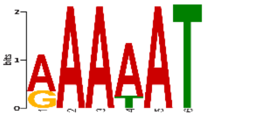 | 7.3e-45 |
| CGTAAATC | 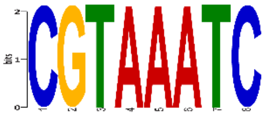 | 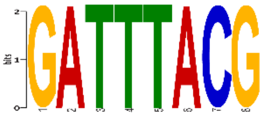 | 1.6e-42 |
| RCGCTCYA | 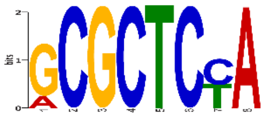 | 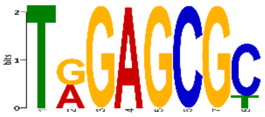 | 4.8e-35 |
| GTMCGCAA | 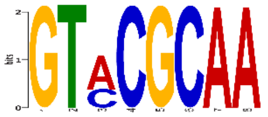 | 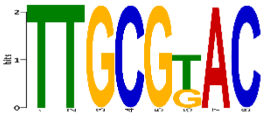 | 8.9e-34 |
| AGMGAAAA | 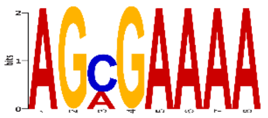 | 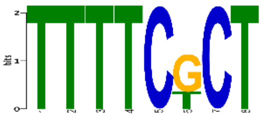 | 3.2e-22 |

**Figure S9** Motif analysis of LSL-1::TY1::EGFP::3XFLAG significant ChIP-seq peaks. List of the 10 most significant motifs enriched in LSL-1::TY1::EGFP::3XFLAG peaks, identified using the MEME-ChIP platform. The most significant motif, [TACBGTA], is very similar to a motif previously described in protein binding microarray studies as an enriched motif in the upstream regions of the germline precursors associated genes (Narasimhan *et al.* 2015). Peaks analyzed here correspond to the merged peaks from two different biological replicates.

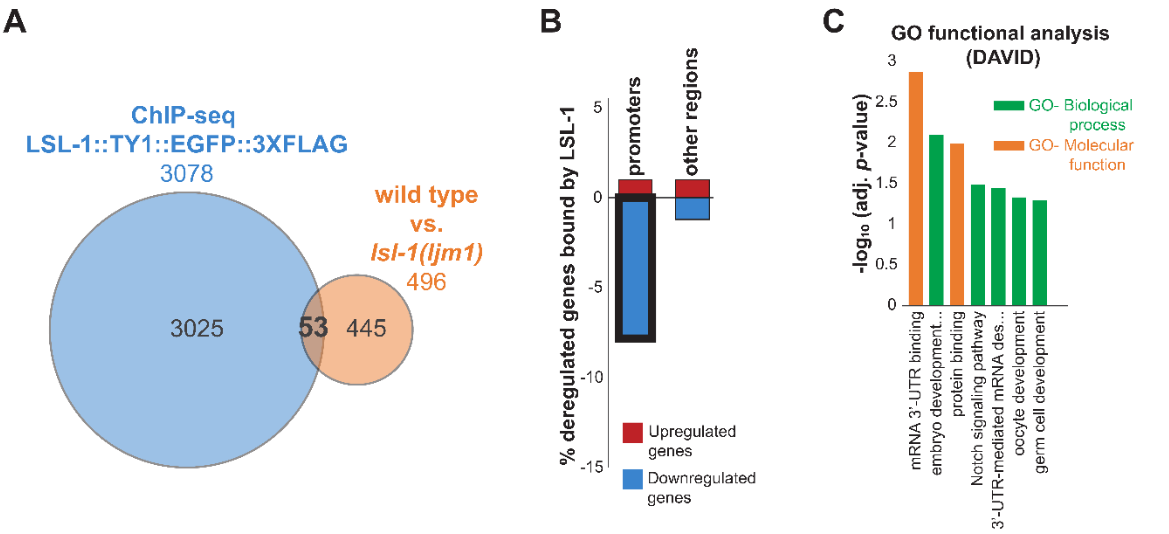


**Figure S10** LSL-1 acts mainly as a transcriptional activator of germline genes. (A) Overlap between LSL-1::TY1::EGFP::3xFLAG ChIP-seq data from modERN resource (Kudron *et al.* 2018) and RNA-seq analysis data of *lsl-1 (ljm1)* DEGs with respect to wild type. Common intercepts are significant DEGs (*q*-value ≤ 0.01, −2 ≥ FC ≥ 2) and significant LSL-1::TY1::EGFP::3xFLAG binding sites (IDR ≤ 0.1%). Overlap was significant (*p*-value ≤ 0.0001). Statistical significance was assessed using cross comparison contingency tables by chi-square test with Yates correction (see Table S2). (B) Graph illustrates the percentage of the 53 significant DEGs in *lsl-1 (ljm1)* animals with respect to wild type and, simultaneously, LSL-1::TY1::EGFP::3xFLAG target genes, distributed by LSL-1 binding site (promoter vs. other regions). Thick line depicts that most significant DEGs were downregulated and directly bound by LSL-1 in their promoter region. (C) Histogram shows DAVID GO term functional analysis (Huang *et al.* 2009) for downregulated genes in *lsl-1(ljm1)* animals with respect to wild type and bound by LSL-1 in their promoter region. Data are plotted as vertical bars that represent the significance of each GO-term. Adj. *p*-values were all significant (adj. *p*-value ≤ 0.05) and correspond to Benjamini–Hochberg correction. Note, symbol (-) in (B) graphic stands for downregulation in the RNA-seq analysis. DEGs, differentially expressed genes; GO, gene ontology.

### SUPPLEMENTAL FIGURES BIBLIOGRAPHY

Angeles-Albores D., R. Y. N. Lee, J. Chan, and P. W. Sternberg, 2016 Tissue enrichment analysis for *C. elegans* genomics. BMC Bioinformatics 17: 366. https://doi.org/10.1186/s12859-016-1229-9

Huang D. W., B. T. Sherman, and R. A. Lempicki, 2009 Bioinformatics enrichment tools: paths toward the comprehensive functional analysis of large gene lists. Nucleic Acids Res. 37: 1–13. https://doi.org/10.1093/nar/gkn923

Kudron M. M., A. Victorsen, L. Gevirtzman, L. W. Hillier, W. W. Fisher, *et al.*, 2018 The ModERN Resource: Genome-Wide Binding Profiles for Hundreds of *Drosophila* and *Caenorhabditis elegans* Transcription Factors. Genetics 208: 937–949. https://doi.org/10.1534/genetics.117.300657

Narasimhan K., S. A. Lambert, A. W. Yang, J. Riddell, S. Mnaimneh, *et al.*, 2015 Mapping and analysis of *Caenorhabditis elegans* transcription factor sequence specificities. eLife 4: e06967. https://doi.org/10.7554/eLife.06967

O'Connor L., J. Gilmour, and C. Bonifer, 2016 The Role of the Ubiquitously Expressed Transcription Factor Sp1 in Tissue-specific Transcriptional Regulation and in Disease. Yale J. Biol. Med. 89: 513–525.

Perkins A., X. Xu, D. R. Higgs, G. P. Patrinos, L. Arnaud, *et al.*, 2016 Krüppeling erythropoiesis: an unexpected broad spectrum of human red blood cell disorders due to KLF1 variants. Blood 127: 1856–1862. https://doi.org/10.1182/blood-2016-01-694331

Storey J. D., and R. Tibshirani, 2003 Statistical significance for genomewide studies. Proc. Natl. Acad. Sci. 100: 9440–9445. https://doi.org/10.1073/pnas.1530509100
