## Supplemental methods for "The zinc-finger transcription factor LSL-1 is a major regulator of the germline transcriptional program in *C. elegans*"

### List of strains used in this study

AV106: *spo-11(ok79) IV/nT1[unc-?(n754) let-?] (IV;V)*

CA1219: *unc-119(ed3) III*; *ieSi21[sun-1p::sun-1::mRuby::sun-1 3'UTR+Cbr-unc-119(+)] IV*

CER414: *pgl-1(cer70[pgl-1::mcherry]) IV*

CWC14: *cep-1(gk138) lsl-1(tm4769) I/hT2[bli-4(e937)let-?(q782)qIs48] (I;III)*

CWC16: *lsl-1(tm4769) I/tmC18[dpy-5(tmls1200)] I; ced-4(n1162) III*

CWC17: *lsl-1(tm4769) I/hT2[bli-4(e937)let-?(q782)qIs48] (I;III); spo-11(ok79) IV/nT1[unc-?(n754)let-?] (IV;V)*

CWC19: *lsl-1(ljm1) I/tmC18[dpy-5(tmls1200)] I*

CWC22: *lsl-1(tm4769) I/tmC18[dpy-5(tmls1200)] I*

CWC39: *lsl-1(ljm1) I/tmC18[dpy-5(tmls1200)]I; ieSi21[sun-1p::sun-1::mRuby::sun-1 3'UTR+Cbr-unc-119(+)] IV*

CWC40: *lsl-1(ljm1) I/tmC18[dpy-5(tmls1200)] I; hpl-2(tm1489) III*

CWC41: *lsl-1(ljm1) I/tmC18[dpy-5(tmls1200)] I; met-2(n4256) III*

CWC44: *lsl-1(tm4769) I/tmC18[dpy-5(tmls1200)] I; hpl-2(tm1489) III*

CWC45: *lsl-1(tm4769) I/tmC18[dpy-5(tmls1200)] I; met-2(n4256) III*

CWC48: *pgl-1(cer70[pgl-1::mcherry]) IV; wgls720[lsl-1::TY1::EGFP::3xFLAG+unc-119(+)] ?*

CWC50: *lsl-1(syb3772[lsl-1::GFP]) I; pgl-1(cer70[pgl-1::mcherry]) IV*

CWC54: *lsl-1(tm4769) I/tmC18[dpy-5(tmls1200)]I;ieSi21[sun-1p::sun-1::mRuby::sun-1 3'UTR+Cbr-unc-119(+)] IV*

CWC59: *lsl-1(ljm1) I/tmC18[dpy-5(tmls1200)] I; met-2(n4256) set-25(n5021) III*

CWC60: *lsl-1(ljm1) I/tmC18[dpy-5(tmls1200)] I; hpl-1(tm1624) X*

CWC67: *lsl-1(tm4769) I/tmC18[dpy-5(tmls1200)] I; hpl-1(tm1624) X*

CWC70: *lsl-1(ljm1) I/tmC18[dpy-5(tmls1200)] I; let-418(n3536) V*

CWC76: *lsl-1(tm4769) I/tmC18[dpy-5(tmls1200)] I; met-2(n4256) set-25(n5021) III*

CWC9: *plk-2(ok1936) lsl-1(tm4769) I/hT2[bli-4(e937)let-?(q782)qIs48] (I;III)*

FR1469: *lsl-1(tm4769) I/hT2[bli-4(e937)let-?(q782)qIs48] (I;III)*

FR1470: *lsl-1(tm4769) I/hT2[bli-4(e937)let-?(q782)qIs48] (I;III); let-418(n3536) V*

FR843: *let-418(n3536) V*

GW638: *met-2(n4256) set-25(n5021) III*

MT13293: *met-2(n4256) III*

MT2547: *ced-4(n1162) III*

OP720: *unc-119(tm4063) III; wgls720[lsl-1::TY1::EGFP::3xFLAG+unc-119(+)] ?*

PFR40: *hpl-2(tm1489) III*

PFR60: *hpl-1(tm1624) X*

PHX3772: *lsl-1(syb3772[lsl-1::GFP]) I*

RB1583: *plk-2(ok1936) I*

VC172: *cep-1(gk138) I*

The loss-of-function allele *lsl-1(ljm1)* of the strain CWC19: *lsl-1(ljm1) I/tmC18[dpy-5(tmls1200)] I* was generated by CRISPR/CAS9 system, using pIK155 (*Peft-3::Cas9::tbb-2 3'UTR*) and AF-ZF-827 oligonucleotide *dpy-10 (cn64)* CRISPR tools described in (Arribere *et al.* 2014 and Katic *et al.* 2015), respectively. Guide RNA was cloned from pMB70, a kind gift from Mike Boxem (Department of Biology Utrecht University; Utrecht, The Netherlands) (Waaijers *et al.* 2013). We inserted two consecutive stop codons at the endogenous *lsl-1* locus, 27 bp downstream the translational initiation site (TIS). The repair template oligonucleotide used was prCW206:

aaaattttattttttacttcagATGTCAATTATTGATGACCGAACGGAT**TG**ATCTTA**A**GACGGCGAGGACTACGAAGCTTCTATAACGgtattttatttcgattctta.

FR1469: *lsl-1(tm4769) I/hT2[bli-4(e937)let-?(q782)qIs48] (I;III)* strain was generated from FX04769: *lsl-1(tm4769) I/(+) I,* which was provided by the National Bioresource Project, *C. elegans* Gene Knockout Consortium; Tokyo, Japan (*C. elegans* Deletion Mutant Consortium 2012).

AV106 strain, *spo-11(ok79) IV/nT1[unc-?(n754) let-?] (IV;V)*, was a kind gift from Anne M. Villeneuve (Stanford University School of Medicine; Stanford CA, USA). CER414 strain, *pgl-1 (cer70[pgl-1::mcherry]) IV*, was kindly provided by Julian Ceron (*C. elegans* Core Facility-IDIBELL; Barcelona, Spain). PHX3772: *lsl-1(syb3772)* is a CRISPR/Cas9 knock-in of GFP sequence at the C-terminal in the endogenous site of *lsl-1* gene, which was generated by SunyBiotech Company (Fuzhou, China). All other strains used in this study were provided by or generated using strains from the *Caenorhabditis* Genetics Center (CGC, University of Minnesota; Minneapolis MN, USA) funded by the National Institutes of Health (NIH; Bethesda MD, USA).

### LSL-1 domains prediction and protein alignments

Protein domains for LSL-1 full-length sequence were predicted by different bioinformatic tools: ScanProsite, available online at https://prosite.expasy.org/scanprosite (de Castro *et al.* 2006); SMART, at https://smart.embl.de (Schultz *et al.* 1998; Letunic *et al.* 2021); and Pfam, at http://pfam.xfam.org (Sonnhammer *et al.* 1996; Mistry *et al.* 2021). LSY-2 was identified as the closest homolog of LSL-1 by BLASTp analysis with *C. elegans* proteins. BLASTp is accessible online at https://blast.ncbi.nlm.nih.gov/Blast.cgi (Altschul *et al.* 1990; Boratyn *et al.* 2012). ZFP57 was identified as the closest human ortholog of LSL-1 by the bioinformatic tools

ALLIANCE, available through the platform https://www.alliancegenome.org (The Alliance of Genome Resources Consortium *et al.* 2020), and PhylomeDB, at http://phylomedb.org (Huerta-Cepas *et al.* 2007), using gene phylogenic trees predictions. Alignments between LSL-1 and its orthologs were performed by BLASTp analysis, and similarity was calculated for the sequence alignments displayed in the Supplemental Material, Figure S2.

### List of antibodies used in this study

Primary antibodies: guinea pig α-HIM-8 1:100 was kindly provided by Abby F. Dernburg (Phillips *et al.* 2005); rabbit α-HTP-3 1:200 was a gift from Monique Zetka (Goodyer *et al.* 2008); and mouse α-Histone H3 Ser10-p (Upstate # 05-866) was used at 1:200.

All secondary antibodies were used at a concentration of 1:200: goat α-rabbit FITC (Jackson ImmunoResearch # 111-095-003); donkey α-guinea pig FITC (Jackson ImmunoResearch # 706-005-148); and goat α-mouse TRITC (Sigma Aldrich # T5393).

### Immunofluorescence in adult hermaphrodite gonads

Worms were dissected (Crittenden *et al.* 1994) in 1x egg buffer [25mM HEPES-NaOH, 118mM NaCl, 48mM KCl, 2mM EDTA and 0.5mM EGTA], 0.1% Tween-20, 20mM $NaN_3$ (EBTA) 24 h post-L4 stage to extrude the gonads and then fixed with 2% formaldehyde for 5 min and flash-frozen on positively charged slides placed on a metal block previously cooled in dry ice. These were then permeabilized by freeze-cracking, postfixed in methanol at −20 °C for 1 min and transferred to 1x phosphate-buffered saline, 0.1% Tween-20 (PBST) at room temperature. Fixed samples were incubated at 4 °C overnight in the primary antibody dilution, washed 3 × 10 min in PBST, and then incubated with the secondary antibody at room temperature for 2 h. Slides were washed 3 × 10 min in PBST, adding 2 μg/ml 2-(4-amidinophenyl)-1H-indole-6-

carboxamidine (DAPI) for 2 min between the second and third washes. Finally, these were mounted with Vectashield H-1000 antifade mounting medium (Vector Laboratories; Burlingame CA, USA), stored at 4 °C, and imaged. To detect endogenous expression of *lsl-1::GFP* and *pgl-1::mCherry*, 1-day adult hermaphrodites were fixed as described above and washed 3 × 10 min in PBST, with DAPI added for 2 min between the second and third washes.

### Germline mitotic region cytological analysis

To analyze the germline mitotic region, at least 20 gonads of each genotype were immunostained using mouse α-Histone H3 Ser10-p antibody (PH3) and DAPI to counterstain DNA. Total number of germ nuclei in the mitotic region was quantified between the distal tip and the transition zone, defining the distal transition zone limit as the first germ cells row where at least two nuclei showed the characteristic crescent shape (Crittenden *et al.* 2006). The mitotic region length was measured in nuclei rows from the distal tip to the transition zone limit, and the mitotic index was determined by the number of PH3-positive nuclei over the total number of germ nuclei in the mitotic region (Maciejowski *et al.* 2006).

### Fluorescence *in situ* hybridization (FISH)

FISH probe hybridization was adapted from (Phillips *et al.* 2009). Briefly, worms were dissected (Crittenden *et al.* 1994) 24 h post-L4 stage to extrude the gonads in EBTA, then fixed with 0.8% ethyleneglycol bis(succinimidylsuccinate) (EGS) in dimethyl formamide for 2 min on positively charged slides, and then incubated in a humid chamber at room temperature for 30 min. Slides were then flash-frozen placed on a metal block previously cooled in dry ice. Next, these were permeabilized by freeze-cracking, postfixed in methanol at −20°C for 1 min, and immediately transfer to 2x saline-sodium citrate, 0.1% Tween-20 (2xSSCT) at room temperature. Slides were incubated in 2x egg buffer, 7.4% formaldehyde (EBF) for 5 min and then rinsed 3 x 5 min in 2xSSCT, adding 50% formamide in 2xSSCT dilution for 5 min between the second and third washes. After the third 2xSSCT wash, slides were incubated in 50% formamide in 2xSSCT dilution at 37 ˚C overnight prior to adding of FISH probe at 37 ˚C. DNA was denatured for 3 min at 95 ˚C, and hybridization was carried out overnight at 37 ˚C in a dark humid chamber. After hybridization, slides were again incubated 1 h in 50% formamide in 2xSSCT dilution at 37 ˚C and washed in 2xSSCT for 10 min, adding 2 µg/ml DAPI for 2 min and finally washing in 2xSSCT for at least 30 min. Slides were then mounted with Vectashield H-1000, stored in darkness at 4 °C, and imaged.

### Acridine orange (AO) staining

Adult hermaphrodite worms, 24-h post-L4-stage, were incubated for 1 h at room temperature and darkness, in plates previously treated with 20µg/ml AO in M9. Then, worms were transferred to clean plates to remove excess of AO, kept in darkness at room temperature for 2 h, and mounted in 2% agar slides. A minimum of 24 gonads per genotype were analyzed, and each scoring experiment was performed with mutant strains and N2 running in parallel. Number of apoptotic corpses per gonad arm was scored using a fluorescence microscope Leica DM1000 LED. Data were pooled from multiple rounds of analyses, and statistical comparation between genotypes was assessed using two-tailed Student's *t*-test with Welch's correction, *p*-value $\leq 0.05$.

### RNA-seq data analysis

The quality of the reads was confirmed with FastQC, retrieved from http://www.bioinformatics.babraham.ac.uk/projects/fastqc/, and the reads were aligned to the *C. elegans* reference genome (version WS220) with TopHat (version 2.1.1)/Bowtie2 (version 2.2.8.0) (Langmead and Salzberg 2012; Kim *et al.* 2013) to obtain the *.bam* files. Read count by gene was obtained by HTSeq-count (Anders *et al.* 2015), and differential gene expression analyses *lsl-1(tm4769)* vs. wild-type, and *lsl-1(ljm1)* vs. wild-type were performed using DESeq2 package (Love *et al.* 2014). Read counts were normalized by estimating size factors, and differential expression was tested against the negative binomial distribution using the Wald test. Multiple test correction was performed via optimized false discovery rate (FDR) approach to obtain an adjusted *p*-value (*q*-value) (Storey and Tibshirani 2003). Genes were defined as differentially expressed genes (DEGs) with a *q*-value ≤ 0.01 (minimum FDR) and −2 ≥ fold change ≥ 2 cutoff. Final lists of significant DEGs for each comparison were converted to Excel sheets; processed lists together with DESeq2 raw data outcome are provided in File S1.

Tissue enrichment analysis for DEGs was performed separately for upregulated and downregulated genes using the Wormbase tool T.E.A. (Angeles-Albores *et al.* 2016). Enriched terms were found significant with an adj. *p*-value ≤ 0.05 obtained from the FDR correction using the Benjamini–Hochberg algorithm; lists of these significant terms were converted to Excel sheets for both *lsl-1(tm4769)* vs. wild-type and *lsl-1(ljm1)* vs. wild-type comparisons and are presented in File S2. Before uploading the DEGs to the T.E.A. resource, genes names were updated to the last Wormbase release at the moment (WS276), using SimpleMine and Gene Name Sanitizer tools available at: https://wormbase.org/tools.

**Chromatin immunoprecipitation (ChIP), data processing, and analysis**

ChIP-seq experiment was conducted by the modERN consortium (Kudron *et al.* 2018) in young adult worms of the *C. elegans* strain OP720, which carry LSL-1::TY1::EGFP::3XFLAG fusion protein (two biological replicates); IP was performed using an anti-GFP antibody.

Sequencing data were aligned to the *C. elegans* reference genome (version WS245) using the Burrows–Wheeler aligner (BWA) (Li and Durbin 2009). Regions significantly enriched in aligned reads were peak-called using ChIP-seq pipeline SPP (Kharchenko *et al.* 2008). Peaks above an irreproducibility discovery rate (IDR) of 0.1% were used to generate the final peaks set (Landt *et al.* 2012). We uploaded the *.bed* optimal IDR thresholded peaks file, available in the modERN website under the accession number ENCFF435YQE, to the Galaxy web platform using the public server at https://usegalaxy.org to analyze the data (Afgan *et al.* 2016). ChIPseeker R/bioconductor package (version 1.18.0) (Yu *et al.* 2015) was used to annotate the *.bed* file. The nearest feature was used with promoter region defined as 2000 bp upstream of a gene, and intergenic region defined as 5000 bp between genes. List of annotated genes bound by LSL-1 was converted to Excel sheets and processed, noncoding RNA genes and genes targeted by high-occupancy target (HOT) regions (black list) (Niu *et al.* 2011) were filtered to obtain the final LSL-1 targeted gene list provided in File S3.

We searched for enriched motifs binding sites of LSL-1 using MEME-ChIP (version 4.11.2) web-based tool (Machanick and Bailey 2011). Input data for MEME-ChIP were composed by central region of the binding sites of 200-bp, thus delimiting this central region as 100 bp around its summit. This range has been reported (Niu *et al.* 2011) to be appropriate for the majority of the enriched motifs for the transcription factors presented in Gerstein *et al.* 2010. The optimal IDR thresholded peaks (IDR 0.1%) obtained from the modERN consortium were chosen for motif discovery. Most significant motifs found by MEME-ChIP were sorted by their e-value, computed by the motif discovery software DREME (Bailey 2011) and MEME (Bailey and Elkan 1994), and presented in Figure S9.

Genome-wide LSL-1 binding profiles were visualized and aligned to the *C. elegans* genome (version WS245) using the Integrated Genome Viewer (IGV) (Thorvaldsdottir *et al.* 2013).

### Functional analysis

Cross comparison of our RNA-seq data and the LSL-1::TY1::EGFP::3XFLAG ChIP-seq analysis showed a significant overlap for both *lsl-1* mutant alleles. Prior to comparing RNA-seq DEGs and ChIP-seq LSL-1 targeted gene lists, gene public names were updated to the last Wormbase release at that time (WS276) using SimpleMine and Gene Name Sanitizer tools available at: https://wormbase.org/tools. Statistical significance was assessed using cross comparison contingency tables by chi-square test with Yates correction, calculated using GraphPad QuickCalcs available at http://www.graphpad.com/quickcalcs/Contingency1.cfm, and presented in Table S2.

Gene ontology (GO) term functional analysis was performed in both *lsl-1* mutant alleles for downregulated genes bound by LSL-1 at their promoter region, using the database for annotation, visualization, and integrated discovery (DAVID version 6.8) NIAID/NIH tool (Huang *et al.* 2009). Significant GO terms ($p$-value ≤ 0.05, unadjusted) are shown, once clustered for most significant categories, in Figure 7B' and Figure S10 C.

### SUPPLEMENTAL METHODS BIBLIOGRAPHY

Afgan E., D. Baker, M. van den Beek, D. Blankenberg, D. Bouvier, *et al.*, 2016 The Galaxy platform for accessible, reproducible and collaborative biomedical analyses: 2016 update. Nucleic Acids Res. 44: W3–W10. https://doi.org/10.1093/nar/gkw343

Altschul S. F., W. Gish, W. Miller, E. W. Myers, and D. J. Lipman, 1990 Basic Local Alignment Search Tool. J. Mol. Biol. 215: 403–410. https://doi.org/10.1016/S0022-2836(05)80360-2

Anders S., P. T. Pyl, and W. Huber, 2015 HTSeq--a Python framework to work with high-throughput sequencing data. Bioinformatics 31: 166–169. https://doi.org/10.1093/bioinformatics/btu638

Angeles-Albores D., R. Y. N. Lee, J. Chan, and P. W. Sternberg, 2016 Tissue enrichment analysis for *C. elegans* genomics. BMC Bioinformatics 17: 366. https://doi.org/10.1186/s12859-016-1229-9

Arribere J. A., R. T. Bell, B. X. H. Fu, K. L. Artiles, P. S. Hartman, *et al.*, 2014 Efficient Marker-Free Recovery of Custom Genetic Modifications with CRISPR/Cas9 in *Caenorhabditis elegans*. Genetics 198: 837–846. https://doi.org/10.1534/genetics.114.169730

Bailey T. L., and C. Elkan, 1994 Fitting a mixture model by expectation maximization to discover motifs in biopolymers. Proc. Int. Conf. Intell. Syst. Mol. Biol. 2: 28–36.

Bailey T. L., 2011 DREME: motif discovery in transcription factor ChIP-seq data. Bioinformatics 27: 1653–1659. https://doi.org/10.1093/bioinformatics/btr261

Boratyn G. M., A. A. Schäffer, R. Agarwala, S. F. Altschul, D. J. Lipman, *et al.*, 2012 Domain enhanced lookup time accelerated BLAST. Biol. Direct 7: 1–12. https://doi.org/10.1186/1745-6150-7-12

*C. elegans* Deletion Mutant Consortium, 2012 Large-Scale Screening for Targeted Knockouts in the *Caenorhabditis elegans* Genome. G3 GenesGenomesGenetics 2: 1415–1425. https://doi.org/10.1534/g3.112.003830

Castro E. de, C. J. A. Sigrist, A. Gattiker, V. Bulliard, P. S. Langendijk-Genevaux, *et al.*, 2006 ScanProsite: detection of PROSITE signature matches and ProRule-associated functional and structural residues in proteins. Nucleic Acids Res. 34: W362–W365. https://doi.org/10.1093/nar/gkl124

Crittenden S. L., E. R. Troemel, T. C. Evans, and J. Kimble, 1994 GLP-1 is localized to the mitotic region of the *C. elegans* germ line. Development 120: 2901–2911.

Crittenden S. L., K. A. Leonhard, D. T. Byrd, and J. Kimble, 2006 Cellular Analyses of the Mitotic Region in the *Caenorhabditis elegans* Adult Germ Line, (J. Schwarzbauer, Ed.). Mol. Biol. Cell 17: 3051–3061. https://doi.org/10.1091/mbc.e06-03-0170

Gerstein M. B., Z. J. Lu, E. L. Van Nostrand, C. Cheng, B. I. Arshinoff, *et al.*, 2010 Integrative Analysis of the *Caenorhabditis elegans* Genome by the modENCODE Project. Science 330: 1775–1787. https://doi.org/10.1126/science.1196914

Goodyer W., S. Kaitna, F. Couteau, J. D. Ward, S. J. Boulton, *et al.*, 2008 HTP-3 Links DSB Formation with Homolog Pairing and Crossing Over during *C. elegans* Meiosis. Dev. Cell 14: 263–274. https://doi.org/10.1016/j.devcel.2007.11.016

Huang D. W., B. T. Sherman, and R. A. Lempicki, 2009 Bioinformatics enrichment tools: paths toward the comprehensive functional analysis of large gene lists. Nucleic Acids Res. 37: 1–13. https://doi.org/10.1093/nar/gkn923

Huerta-Cepas J., H. Dopazo, J. Dopazo, and T. Gabaldón, 2007 The human phylome. Genome Biol. 8: R109. https://doi.org/10.1186/gb-2007-8-6-r109

Katic I., L. Xu, and R. Ciosk, 2015 CRISPR/Cas9 Genome Editing in *Caenorhabditis elegans* : Evaluation of Templates for Homology-Mediated Repair and Knock-Ins by Homology-Independent DNA Repair. G3 GenesGenomesGenetics 5: 1649–1656. https://doi.org/10.1534/g3.115.019273

Kharchenko P. V., M. Y. Tolstorukov, and P. J. Park, 2008 Design and analysis of ChIP-seq experiments for DNA-binding proteins. Nat. Biotechnol. 26: 1351–1359. https://doi.org/10.1038/nbt.1508

Kim D., G. Pertea, C. Trapnell, H. Pimentel, R. Kelley, *et al.*, 2013 TopHat2: accurate alignment of transcriptomes in the presence of insertions, deletions and gene fusions. Genome Biol. 14: R36. https://doi.org/10.1186/gb-2013-14-4-r36

Kudron M. M., A. Victorsen, L. Gevirtzman, L. W. Hillier, W. W. Fisher, *et al.*, 2018 The ModERN Resource: Genome-Wide Binding Profiles for Hundreds of *Drosophila* and *Caenorhabditis elegans* Transcription Factors. Genetics 208: 937–949. https://doi.org/10.1534/genetics.117.300657

Landt S. G., G. K. Marinov, A. Kundaje, P. Kheradpour, F. Pauli, *et al.*, 2012 ChIP-seq guidelines and practices of the ENCODE and modENCODE consortia. Genome Res. 22: 1813–1831. https://doi.org/10.1101/gr.136184.111

Langmead B., and S. L. Salzberg, 2012 Fast gapped-read alignment with Bowtie 2. Nat. Methods 9: 357–359. https://doi.org/10.1038/nmeth.1923

Letunic I., S. Khedkar, and P. Bork, 2021 SMART: recent updates, new developments and status in 2020. Nucleic Acids Res. 49: D458–D460. https://doi.org/10.1093/nar/gkaa937

Li H., and R. Durbin, 2009 Fast and accurate short read alignment with Burrows-Wheeler transform. Bioinformatics 25: 1754–1760. https://doi.org/10.1093/bioinformatics/btp324

Love M. I., W. Huber, and S. Anders, 2014 Moderated estimation of fold change and dispersion for RNA-seq data with DESeq2. Genome Biol. 15: 550. https://doi.org/10.1186/s13059-014-0550-8

Machanick P., and T. L. Bailey, 2011 MEME-ChIP: motif analysis of large DNA datasets. Bioinformatics 27: 1696–1697. https://doi.org/10.1093/bioinformatics/btr189

Maciejowski J., N. Ugel, B. Mishra, M. Isopi, and E. J. A. Hubbard, 2006 Quantitative analysis of germline mitosis in adult *C. elegans*. Dev. Biol. 292: 142–151. https://doi.org/10.1016/j.ydbio.2005.12.046

Mistry J., S. Chuguransky, L. Williams, M. Qureshi, G. A. Salazar, *et al.*, 2021 Pfam: The protein families database in 2021. Nucleic Acids Res. 49: D412–D419. https://doi.org/10.1093/nar/gkaa913

Niu W., Z. J. Lu, M. Zhong, M. Sarov, J. I. Murray, *et al.*, 2011 Diverse transcription factor binding features revealed by genome-wide ChIP-seq in *C. elegans*. Genome Res. 21: 245–254. https://doi.org/10.1101/gr.114587.110

Phillips C. M., C. Wong, N. Bhalla, P. M. Carlton, P. Weiser, *et al.*, 2005 HIM-8 Binds to the X Chromosome Pairing Center and Mediates Chromosome-Specific Meiotic Synapsis. Cell 123: 1051–1063. https://doi.org/10.1016/j.cell.2005.09.035

Phillips C. M., K. L. McDonald, and A. F. Dernburg, 2009 Cytological Analysis of Meiosis in *Caenorhabditis elegans*, pp. 171–195 in *Meiosis*, edited by Keeney S. Humana Press, Totowa, NJ.

Schultz J., F. Milpetz, P. Bork, and C. P. Ponting, 1998 SMART, a simple modular architecture research tool: Identification of signaling domains. Proc. Natl. Acad. Sci. 95: 5857–5864. https://doi.org/10.1073/pnas.95.11.5857

Sonnhammer E. L. L., S. R. Eddy, and R. Durbin, 1996 Pfam: A comprehensive database of protein domain families based on seed alignments. PROTEINS Struct. Funct. Genet. 28: 405–420. https://doi.org/10.1002/(sici)1097-0134(199707)28:3<405::aid-prot10>3.0.co;2-l

Storey J. D., and R. Tibshirani, 2003 Statistical significance for genomewide studies. Proc. Natl. Acad. Sci. 100: 9440–9445. https://doi.org/10.1073/pnas.1530509100

The Alliance of Genome Resources Consortium, J. Agapite, L.-P. Albou, S. Aleksander, J. Argasinska, *et al.*, 2020 Alliance of Genome Resources Portal: unified model organism research platform. Nucleic Acids Res. 48: D650–D658. https://doi.org/10.1093/nar/gkz813

Thorvaldsdottir H., J. T. Robinson, and J. P. Mesirov, 2013 Integrative Genomics Viewer (IGV): high-performance genomics data visualization and exploration. Brief. Bioinform. 14: 178–192. https://doi.org/10.1093/bib/bbs017

Waaijers S., V. Portegijs, J. Kerver, B. B. L. G. Lemmens, M. Tijsterman, *et al.*, 2013 CRISPR/Cas9-Targeted Mutagenesis in *Caenorhabditis elegans*. Genetics 195: 1187–1191. https://doi.org/10.1534/genetics.113.156299

Yu G., L.-G. Wang, and Q.-Y. He, 2015 ChIPseeker: an R/Bioconductor package for ChIP peak annotation, comparison and visualization. Bioinformatics 31: 2382–2383. https://doi.org/10.1093/bioinformatics/btv145
