## Supplemental tables for "The zinc-finger transcription factor LSL-1 is a major regulator of the germline transcriptional program in *C. elegans*"

This PDF file includes:

Tables S1 and S2

Table S1 Brood size, survival rate, and incidence of males (22 °C and 25 °C)

| Genotype | Mean brood size[a] | Viability (%) | Incidence of males (%) | $n$[b] |
|---|---|---|---|---|
| wild type$^{22\,°C}$ | 241.33 ± 46.45 | 98.62 | 0.07 | 6 |
| *lsl-1(tm4769)*$^{22\,°C}$ | 19.92 ± 19.33*** | 0.00 | n/a | 12 |
| *lsl-1(ljm1)*$^{22\,°C}$ | 8.75 ± 20.42*** | 22.86 | 8.33 | 12 |
| wild type$^{25\,°C}$ | 196.43 ± 42.37 | 94.46 | 0.09 | 46 |
| *lsl-1(tm4769)*$^{25\,°C}$ | 0.05 ± 0.22*** | 0.00 | n/a | 40 |
| *lsl-1(ljm1)*$^{25\,°C}$ | 0.13 ± 0.41*** | 20.00 | 0.00 | 38 |

[a] Data correspond to the mean ± SD of the total number of eggs laid per hermaphrodite parent. Statistical comparison between wild type and each genotype performed by two-tailed Student's *t*-test with Welch's correction. *** $p$-value ≤ 0.001.

[b] Total number of parental hermaphrodites per genotype.

n/a not applicable

Table S2 Cross-comparison contingency tables summary

| RNA-seq sample | DEGs | LSL-1 binding sites (*n* = 3078) | *p*-value |
|---|---|---|---|
| *lsl-1(tm4769) vs* wild type | 2078 | 388 | 0.0001*** |
| *lsl-1(ljm1) vs* wild type | 496 | 53 | 0.0061** |

DEGs, differentially expressed genes; ****p*-value ≤ 0.001, ***p*-value ≤ 0.01, by chi-square test with Yates correction.
